# Complex genomes of early nucleocytoviruses revealed by ancient origins of viral aminoacyl-tRNA synthetases

**DOI:** 10.1101/2023.12.10.570958

**Authors:** Soichiro Kijima, Hiroyuki Hikida, Tom O. Delmont, Morgan Gaïa, Hiroyuki Ogata

## Abstract

Aminoacyl-tRNA synthetases (aaRSs), also known as tRNA ligases, are essential enzymes in translation. Owing to their functional essentiality, these enzymes are conserved in all domains of life and used as informative markers to trace the evolutionary history of cellular organisms. Unlike cellular organisms, viruses generally lack aaRSs because of their obligate parasitic nature, but several large and giant DNA viruses in the phylum *Nucleocytoviricota* encode aaRSs in their genomes. The discovery of viral aaRSs led to the idea that the phylogenetic analysis of aaRSs can shed light on ancient viral evolution. However, conflicting results have been reported from previous phylogenetic studies: one posited that nucleocytoviruses recently acquired their aaRSs from their host eukaryotes, while another hypothesized that the viral aaRSs have ancient origins. Here, we investigated 4,168 nucleocytovirus genomes, including metagenome-assembled genomes derived from large-scale metagenomic studies. In total, we identified 780 viral aaRS sequences in 273 viral genomes. We generated and examined phylogenetic trees of these aaRSs with a large set of cellular sequences to trace evolutionary relationships between viral and cellular aaRSs. The analyses firmly establish that the origins of some viral aaRSs predate the last common eukaryotic ancestor. Inside viral aaRS clades, we identify intricate evolutionary trajectories of viral aaRSs with horizontal transfers, losses, and displacements. Overall, these results suggest that ancestral nucleocytoviruses already developed complex genomes with an expanded set of aaRSs in the proto-eukaryotic era.

## Introduction

*Nucleocytoviricota* is a phylum of dsDNA viruses (nucleocytoviruses), formerly known as nucleo-cytoplasmic large DNA viruses, which infect diverse eukaryotes from microeukaryotes to animals [1–3]. These viruses have large genomes that can exceed 1 Mbp, with over 1,000 genes [4,5]. Their genomes encode genes that were previously considered to be exclusive to cellular organisms, such as those related to energy production, chromatin remodeling, and translation [4,6–9]. Although these cellular hallmark genes in viral genomes were speculated to be derived from cellular organisms, their origin and evolution remain largely elusive. Some proposed the fourth-domain hypothesis based on deep branches of these virus-encoded cellular hallmark genes independent from those encoded in the three established domains of life [4,10–12]. Under this rather provocative hypothesis, nucleocytoviruses originated from ancient and extinct cellular organisms with a complete set of cellular genes and experienced reductive evolution, thereby still retaining many cellular hallmark genes. Meanwhile, others proposed an alternative model in which a small viral ancestor accumulated genes from cellular organisms, like a “gene robber,” by horizontal gene transfer (HGT) or *de novo* gene creation [13–17]. All scenarios, however, suppose that nucleocytoviruses and their host organisms underwent long-lasting interactions. The phylogeny of genes conserved within *Nucleocytoviricota* supported the occurrence of such interactions by demonstrating that ancestral nucleocytoviruses had already appeared in the proto-eukaryotic era, a period before the last eukaryotic common ancestor (LECA) during which the complex and unique traits of eukaryotic cells were developed [3,18,19]. Tight evolutionary associations between eukaryotes and nucleocytoviruses are also supported by HGTs, some of which date back to the proto-eukaryotic period (e.g., two largest subunits of the viral DNA-dependent RNA polymerase, actin-related protein, topoisomerase type IIA) [3,20–22].

Aminoacyl-tRNA synthetases (aaRSs) are among the cellular hallmark enzymes encoded by nucleocytoviruses whose evolution has been controversial [23]. These enzymes have a central role in decoding genetic codes and are conserved in all three domains of life, but small viruses rarely encode them. A nucleocytovirus, Acanthamoeba polyphaga mimivirus (APMV), is the first virus found to encode aaRSs (ArgRS, CysRS, MetRS, and TyrRS) [4]. Isolation of its relative (megavirus chilensis) further expanded the repertoire of viral aaRSs from four to seven (IleRS, TrpRS, and AsnRS, in addition to the set encoded by APMV) [24]. Discoveries of nucleocytoviruses with an increasing number of aaRSs raised expectations that their ancestor had a complete set of 20 aaRSs, which would support the reductive evolution scenario [23]. However, phylogenetic analysis of these aaRSs suggested their recent origins by gradual accumulation through HGT from eukaryotic hosts based on polyphyly and tree topologies suggesting eukaryotic origins for some aaRSs [14]. Recently, nucleocytoviruses with more enriched sets of aaRSs were discovered, but the uncertainty remained. Metagenome-derived klosneuvirus was found to encode 19 aaRSs and their phylogenies supported scenarios of recent origins for most of these genes; 17 out of 19 genes appeared to have been horizontally transferred from different eukaryotes [25]. In contrast, a study of tupanviruses, which were isolated from extreme environments and found to encode the complete set of 20 aaRSs, claimed that the origins of tupanvirus aaRSs are not necessarily explained by recent HGT from eukaryotes [26].

aaRSs are conserved in all domains of life, and therefore they are useful markers to resolve ancient evolution of cellular organisms [27]. However, unlike other marker genes to resolve the tree of life, evolutionary trajectories of aaRSs are complex with frequent HGT and subsequent displacements [28,29]. Eukaryotic aaRSs show particularly complex evolutionary histories due to multiple sets of homologs that function in different translational compartments: cytosol, mitochondria, and chloroplasts (plastids). These characteristics might have hampered an unequivocal interpretation of phylogenetic trees of viral aaRSs. Furthermore, previous phylogenetic analyses of viral aaRSs used a small number of sequences from nucleocytoviruses and incomprehensive taxon sampling of cellular organisms [14,25,26]. Thus, the ability to resolve their evolutionary histories could have been limited by computational artifacts (e.g., long branch attraction). In the present study, we revisit the origins of nucleocytovirus aaRSs by using recently reconstructed metagenome-assembled genomes (MAGs) [30–32] as well as cellular aaRSs from a broad range of taxa [33]. By reconstructing large-scale phylogenetic trees of aaRSs, we reliably identified the origins of some viral aaRSs, which highlight the ancient genome complexity of nucleocytoviruses.

## Results

### Identification of nucleocytovirus aaRSs

From 224 reference nucleocytovirus genomes and 3,578 non-redundant nucleocytovirus MAGs (Table S1), we identified 780 aaRS sequences in 273 genomes (Table S2). PheRS, GlyRS, and LysRS have multiple types of enzymes, but those detected in nucleocytoviruses were all related to a single type: the cellular PheRS ɑ subunit, GlyRS-1, and LysRS-II, respectively. The most abundant aaRSs were associated with AT-rich codons, namely 144 AsnRSs (codons: AAU/C), 125 IleRSs (AUU/C/A), and 84 TyrRSs (UAU/C), representing 18%, 16%, and 11% of the total aaRSs, respectively (Fig. S1). Out of 780 aaRSs, 730 (93.6%) were encoded by viruses in the order *Imitervirales* (Fig. 1, Table S2). Within this order, an internal monophyletic clade consisting of 64 genomes harbored 398 aaRSs, which is over half of the aaRSs encoded in this order. This clade included nucleocytoviruses known to encode an expanded set of aaRSs, such as mimiviruses, tupanviruses, and klosneuviruses [4,25,26] (Supplementary Data 1).

**Figure 1.**
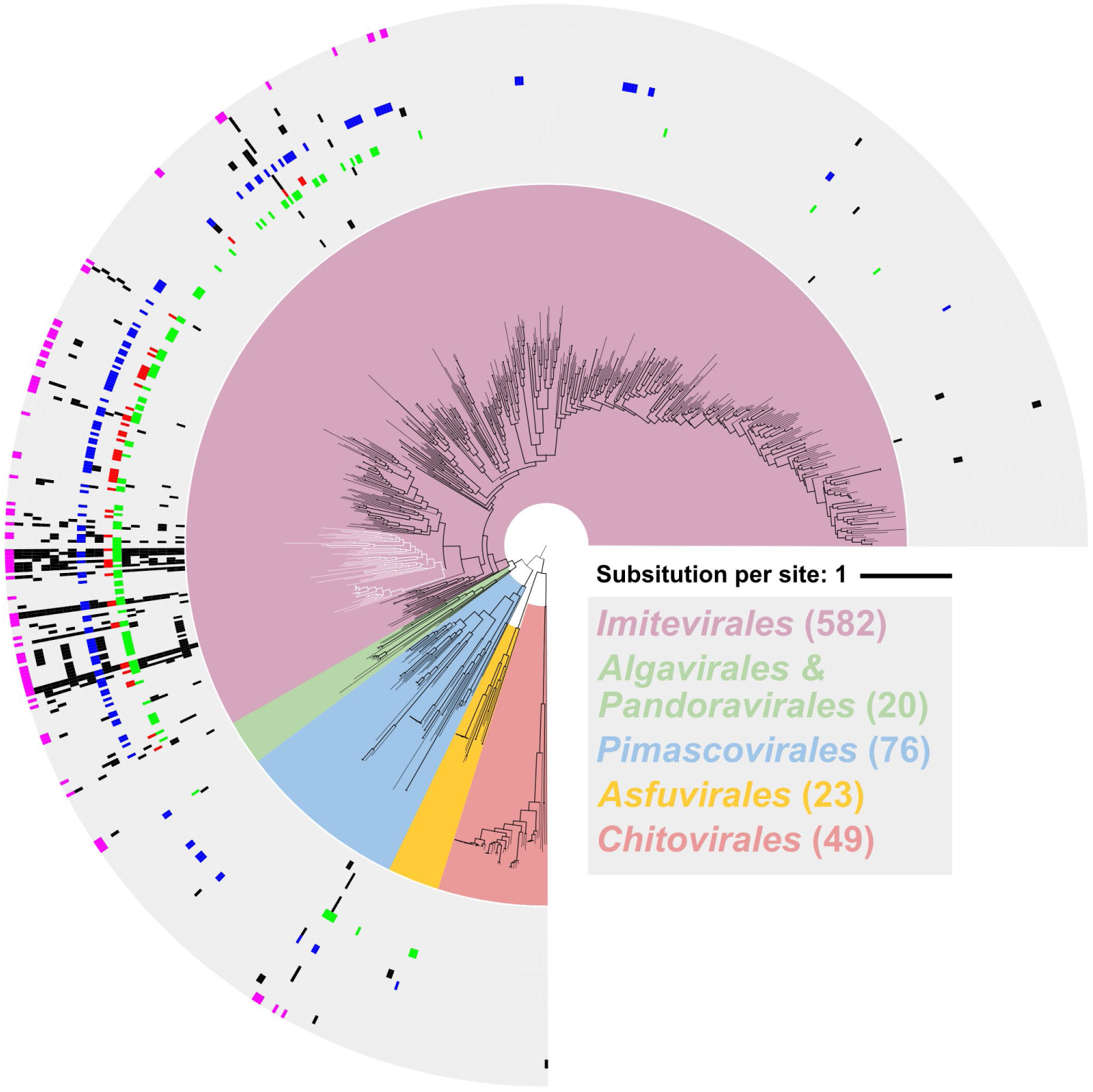
A phylogenetic tree of nucleocytovirus species based on the concatenated sequences of eight core genes. Colors in the tree background represent each viral order. As the sequences from Algavirales and Pandoravirales were limited, they are represented with the same color. White branches indicate the aaRS-rich clade designated in the main text. Branches with thick lines were significantly supported (≥80% SH-aLRT & ≥95% UFB). The layers surrounding the tree represent the distribution of aaRSs, alphabetically ordered by single-character amino acid codes, from inner to outer layer. IleRS, LysRS, AsnRS, and TyrRS are highlighted in green, red, blue, and magenta, respectively. The substitution model was Q.pfam+F+R10. Asfuvirales and Chitovirales were used as outgroups following the method reported by Aylward et al. [1].

### Evolution of nucleocytovirus aaRSs

To elucidate the evolutionary histories of nucleocytovirus aaRSs, we constructed their phylogenetic trees together with a comprehensive set of cellular sequences from a previous study [33]. This dataset covers wide taxonomic groups: 64 eukaryotic species from 26 phyla, 142 bacterial species from 54 phyla, and 76 archaeal species from 11 phyla. The eukaryotic sequences encompassed both cytosolic and organellar sequences. We estimated their evolutionary histories based on tree topology and their statistical support: ultra-fast bootstrap (UFB; >95%) and Shimodaira–Hasegawa approximate likelihood ratio test (SH-aLRT; >80%). We further constructed subsection trees for some aaRSs to improve the resolution of tree topologies. We assigned an evolutionary scenario for each nucleocytovirus aaRS clade, as shown in Table 1. The scenarios were mainly separated into six categories: (1) ancient HGT between proto-eukaryotes and nucleocytoviruses (proto-Euk&V); (2) ancient HGT between eukaryotes and nucleocytoviruses, but not necessarily dating back to proto-eukaryotes (ancient Euk&V); (3) recent HGT from eukaryotes to nucleocytoviruses (recent Euk&V); (4) HGT from nucleocytoviruses to eukaryotes (V to Euk); (5) HGT between eukaryotes and nucleocytoviruses for which the timing of transfer is less clear (other Euk&V); and (6) HGT between prokaryotes and nucleocytoviruses (Prok&V). We describe details of selected notable cases below, while others are presented in Supplementary Information.

**Table 1.**
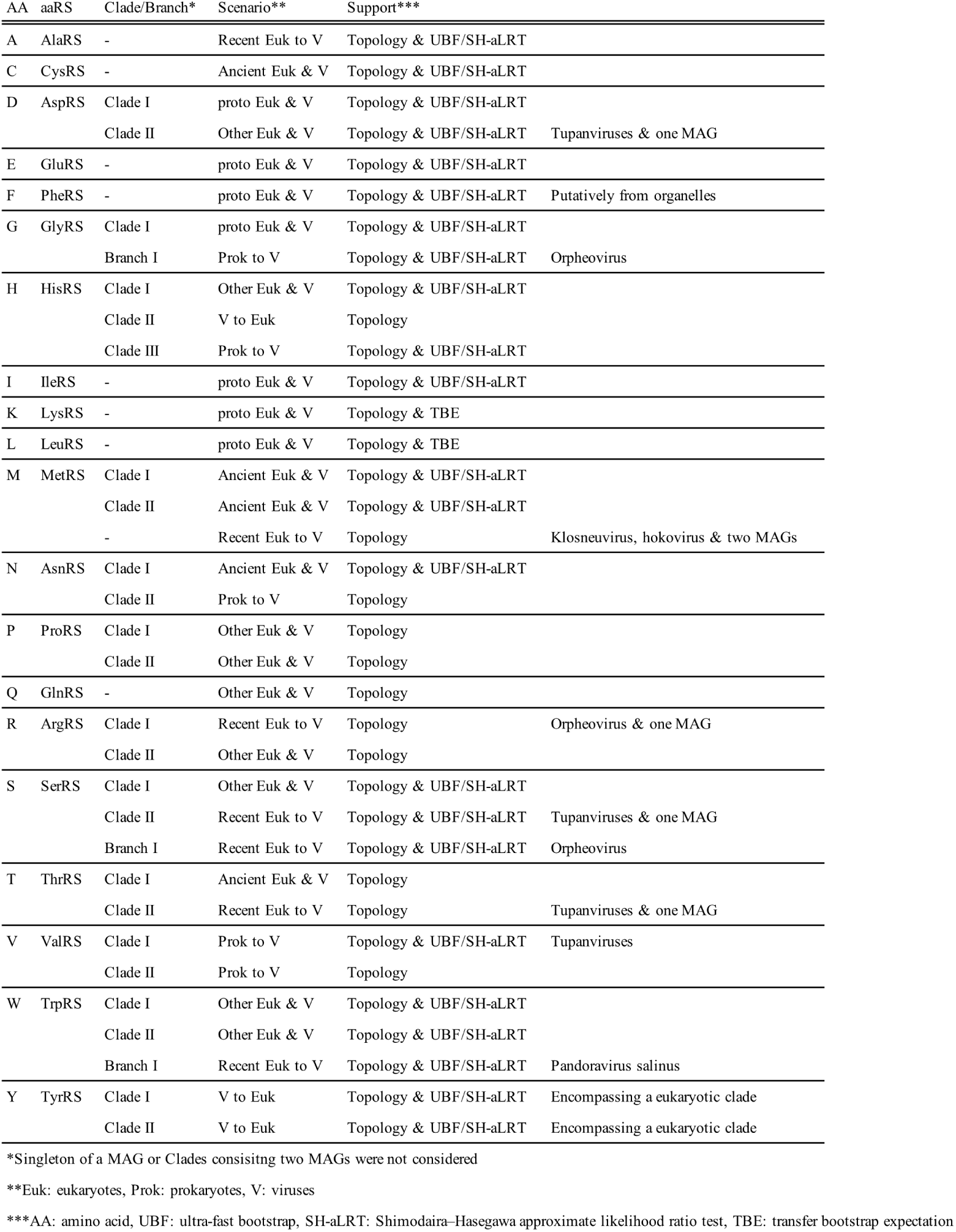
Scenarios for each aaRS.

### Ancient HGT between proto-eukaryotes and nucleocytoviruses (proto-Euk&V)

We identified HGTs between nucleocytoviruses and proto-eukaryotes in GlyRS, IleRS, AspRS, GluRS, LysRS, and LeuRS (Figs. 2 and S2). We describe details of GlyRS and IleRS below, and AspRS, GluRS, LysRS, and LeuRS in Supplementary Information.

**Figure 2.**
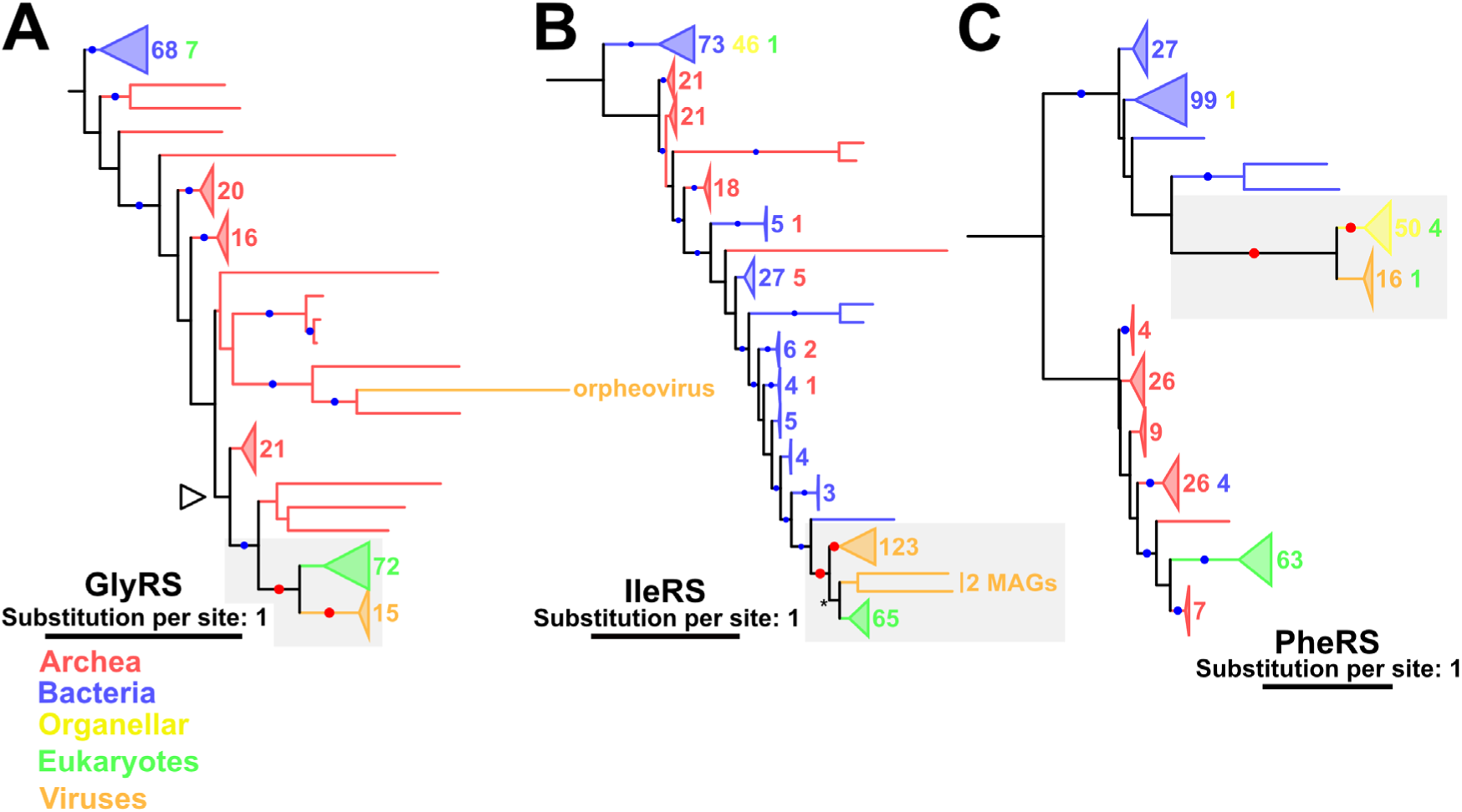
Phylogenetic trees of (A) GlyRS, (B) IleRS, and (C) PheRS. Red and blue dots indicate significant statistical support (≥80% SH-aLRT & ≥95% UFB). The numbers next to the collapsed clades and their colors indicate the number of sequences and their sources, respectively. Red dots and gray area indicate the nodes and the phylogenetic relationships described in the main text, respectively. (A) An empty arrowhead indicates the node from which subsection trees were constructed. (B) An asterisk indicates the node supported by UFB = 94% and SH-aLRT = 96.8%. The substitution models were (A) LG+R9, (B) Q.pfam+F+R10, and (C) LG+R9. The root was decided following the work of Furukawa et al. [33].

In the tree of GlyRS, three domains were largely separated, with a major eukaryotic clade branching from within the archaea domain (Fig. 2A). Two small eukaryotic clades were also found in bacterial clades, but species encoding these GlyRSs were also represented in the major eukaryotic clade. Therefore, the small eukaryotic clades likely consist of organellar GlyRSs derived from bacteria, while the root of the major eukaryotic clade corresponds to LECA. A majority of nucleocytovirus GlyRS sequences formed the sister clade to the major eukaryotic clade. Grouping of the viral GlyRSs with eukaryotic ones and the monophyly of the viral GlyRSs were statistically supported. However, the monophyly of the eukaryotic clade was not supported. We hence generated a subsection tree around the viral and eukaryotic GlyRSs, which supported the monophyly of the eukaryotic GlyRSs (Fig. S3A). These results indicate an ancient HGT for GlyRS between nucleocytoviruses and proto-eukaryotes.

A similar scenario was inferred for IleRS. In the IleRS tree (Fig. 2B), a monophyletic group of all viral sequences and most eukaryotic sequences except one in a bacterial clade was confirmed. Furthermore, a large majority of viral sequences formed a supported monophyletic group, while the monophyly of the clade containing eukaryotic sequences was not supported, which hampers the unambiguous placement of LECA. However, the clade of the eukaryotic sequence and two viral MAGs sister to the eukaryotic group had relatively high support values (UFB = 94%, SH-aLRT = 96.8%), although the UFB value did not meet the threshold that we applied (UFB > 95%). This led us to assume that the root of this clade corresponds to the time of LECA or earlier. Therefore, we conclude that viral IleRSs experienced an ancient HGT between nucleocytoviruses and proto-eukaryotes.

We also identified a putative and ancient HGT between organelles and nucleocytoviruses for PheRS. In the PheRS tree, bacterial and archaeal sequences are largely separated (Fig. 2C). Eukaryotic (nuclear) PheRSs were grouped with archaeal sequences, while nucleocytovirus PheRSs were grouped with bacterial and organellar sequences, indicating different origins for these PheRSs. This organellar clade included all organellar sequences except one and covered a broad range of eukaryotic taxa, suggesting that its root corresponds to the time of LECA. The statistically supported monophyly of organellar sequences and the grouping of the organellar and viral sequences suggest an ancient HGT between viruses and proto-eukaryotes (or organelles), which occurred before LECA.

### Ancient HGT between eukaryotes and nucleocytoviruses (ancient Euk&V)

We identified other cases of ancient HGTs between eukaryotes and nucleocytoviruses. In these cases, the HGT events were inferred to have occurred before the radiation of several major lineages of eukaryotes, although it was unclear whether these events date back to the proto-eukaryotic era (i.e., before LECA). The trees of MetRS, CysRS, and AsnRS suggested this scenario. A similar case was observed for ThrRS, although the scenario was not statistically supported for it (Fig. S4). We here describe the cases for MetRS and CysRS.

The MetRS tree contains two phylogenetically distant large eukaryotic clades (E-clades I and II) (Fig. 3A); one of these clades (E-clade II) contains four viral sequences. Each of the clades is sister to a clade consisting of viral MetRSs (V-clades I and II). In both clades, the monophyly of eukaryotes, the monophyly of nucleocytoviruses, and the grouping of these two clades were statistically supported, indicating the occurrence of HGTs between eukaryotes and nucleocytoviruses before the divergence of the respective eukaryotic clades. Members of the two eukaryotic clades did not overlap: E-clade I mainly included species of the Amorphea and Archaeplastida supergroups and E-clade II included members of the SAR supergroup (Supplementary Data 2). This phylogenetic distribution indicates that eukaryotes acquired their MetRS twice. As the two eukaryotic clades consist of different supergroups, it was unclear whether the roots of these eukaryotic clades correspond to LECA. Nevertheless, the HGTs between eukaryotes and viruses occurred before the divergence of each eukaryotic clade. Thus, we classified the HGTs of MetRS between E-clades and V-clades as ancient events. A clade of viral AsnRSs (Clade I) also suggested a similar evolutionary history (Fig. 3B, Supplementary Information).

**Figure 3.**
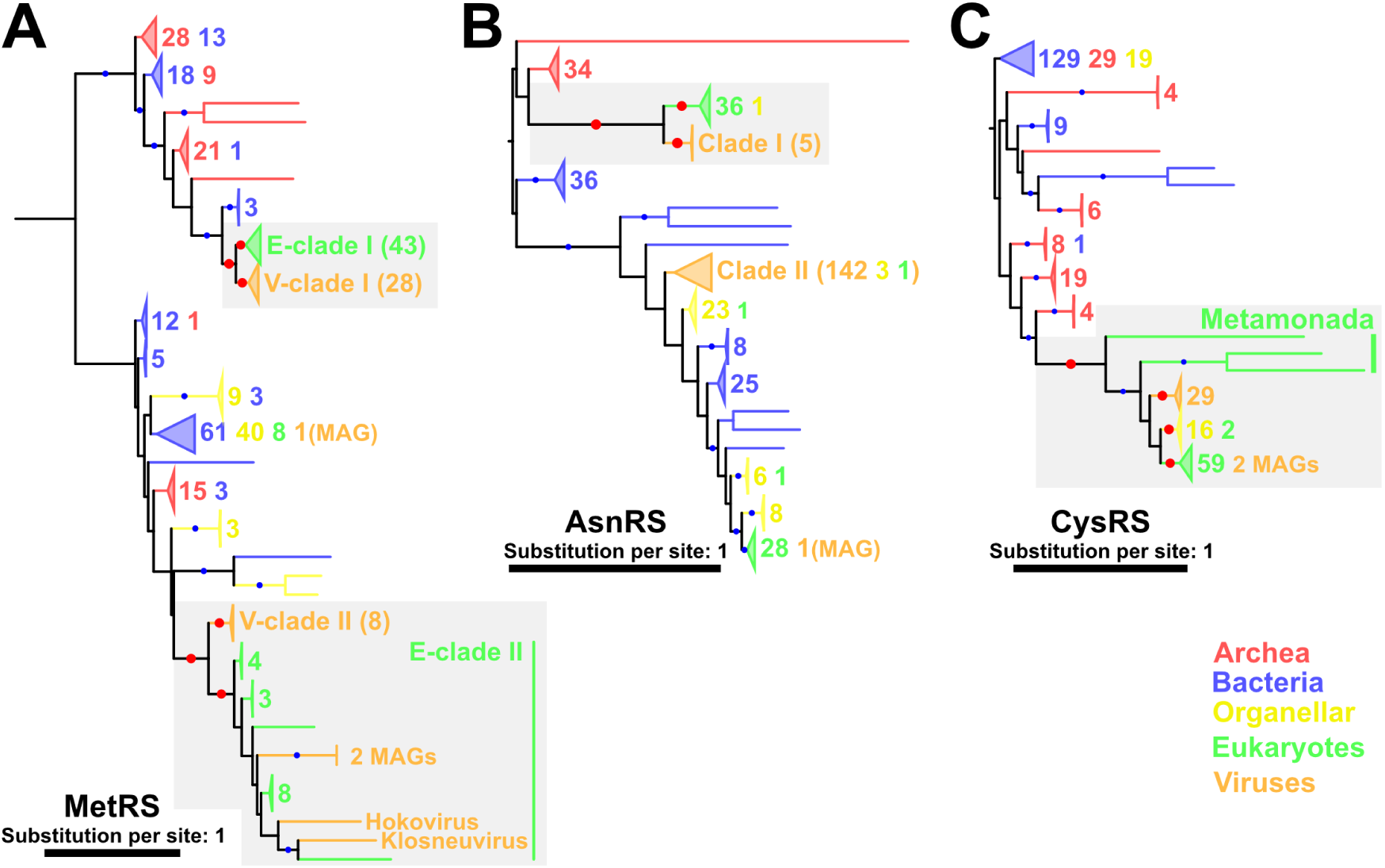
Phylogenetic trees of (A) MetRA, (B) AsnRS, and (C) CysRS. Red and blue dots indicate significant statistical support (≥80% SH-aLRT & ≥95% UFB). The numbers next to the collapsed clades and their colors indicate the number of sequences and their sources, respectively. Some clades are named and the numbers in brackets indicate the number of sequences included within them. Red dots and gray area indicate the nodes and the phylogenetic relationships described in the main text, respectively. The substitution models were (A) LG+R9, (B) Q.pfam+F+R10, and (C) LG+R9. The root was decided following the work of Furukawa et al. [33].

The CysRS tree depicts another scenario (Fig. 3C). This tree has a statistically supported clade consisting of nucleocytoviral, eukaryotic, and organellar sequences. The monophyly of nucleocytovirus sequences and the monophyly of organellar sequences were also statistically supported. Most eukaryotic sequences also formed a statistically supported clade. Only sequences from Metamonada species (*Trichomonas vaginalis*, *Spironucleus salmonicida*, and *Giardia intestinalis*) were located outside of the group of viral, organellar, and other eukaryotic sequences. These results suggest an ancient HGT for CysRS between nucleocytoviruses and eukaryotes before the divergence of most eukaryote lineages.

### Recent HGT from eukaryotes to nucleocytoviruses (Recent Euk to V)

In addition to the ancient HGTs between eukaryotes and nucleocytoviruses, we identified recent transfers from eukaryotes to nucleocytoviruses in AlaRS, TrpRS, ArgRS, SerRS, MetRS, and ThrRS (Figs. 3, 4, S4, and S5). Here, we mainly describe the trees of AlaRS and TrpRS.

**Figure 4.**
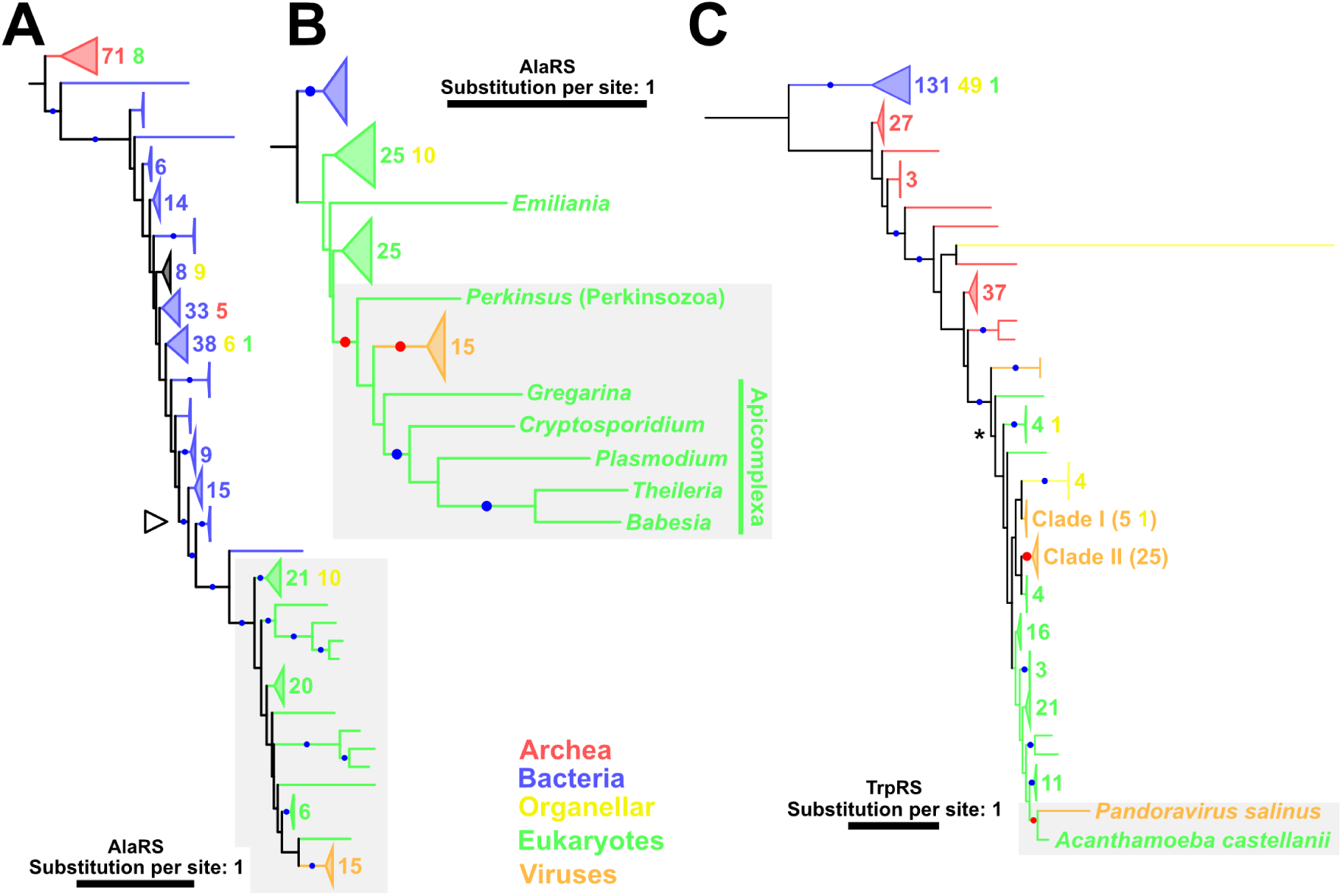
(A) A phylogenetic tree and (B) its subsection tree for AlaRS and (C) a phylogenetic tree of TrpRS. Red and blue dots indicate significant statistical support (≥80% SH-aLRT & ≥95% UFB). The numbers next to the collapsed clades and their colors indicate the number of sequences and their sources, respectively. Red dots and gray area indicate the nodes and the phylogenetic relationship described in the main text, respectively. (A) An empty arrowhead indicates the node from which subsection trees were constructed. (B) For eukaryotic sequences, only names of genera are shown. The substitution models were (A) LG+R10, (B) LG+I+I+R7, and (C) LG+R9. The root was decided following the work of Furukawa et al. [33].

In the AlaRS tree, nucleocytovirus AlaRSs formed a statistically supported monophyletic group (Fig. 4A). This viral AlaRS clade forms a statistically supported clade together with eukaryotic AlaRSs, but the position of the viral clade within eukaryotic branches was unclear. To infer the position of the viral clade, we constructed a subsection tree around the viral and eukaryotic clade (Fig. 4B). The viral AlaRSs still formed a statistically supported clade, which was grouped together with AlaRSs from Apicomplexa and Perkinsozoa (two eukaryotic phyla belonging to Alveolata). This grouping of two eukaryotic phyla and a viral clade was statistically supported (Fig. 4B). Because Perkinsozoa and Apicomplexa are closely related phyla within Alveolata [34], this clade likely represents the vertical evolution of these eukaryotes. Taken together, we conclude that an HGT occurred for AlaRS from eukaryotes to nucleocytoviruses during the divergence of these eukaryotic phyla.

Several aaRSs identified in the genomes of isolated viruses also showed clear evidence of recent transfer from eukaryotes. One example was found in TrpRS encoded by pandoravirus salinus (Fig. 4C). This TrpRS was grouped together with that from *Acanthamoeba castellanii* with statistical support. Because *A. castellanii* is one of the host organisms of the pandoravirus, this phylogenetic relationship suggests that this viral TrpRS was acquired from its host by HGT. Similar cases were suggested for an ArgRS encoded by orpheovirus and SerRSs encoded by tupanvirus and orpheovirus, although the sources of these HGTs were unclear (Fig. S5, Supplementary Information).

### HGT from nucleocytoviruses to eukaryotes (V to Euk)

Putative HGTs from viruses to eukaryotes were suggested for TyrRS, SerRS, AsnRS, and HisRS (Figs. 5, S5, and S6). A case found in the HisRS tree is described in Supplementary Information.

**Figure 5.**
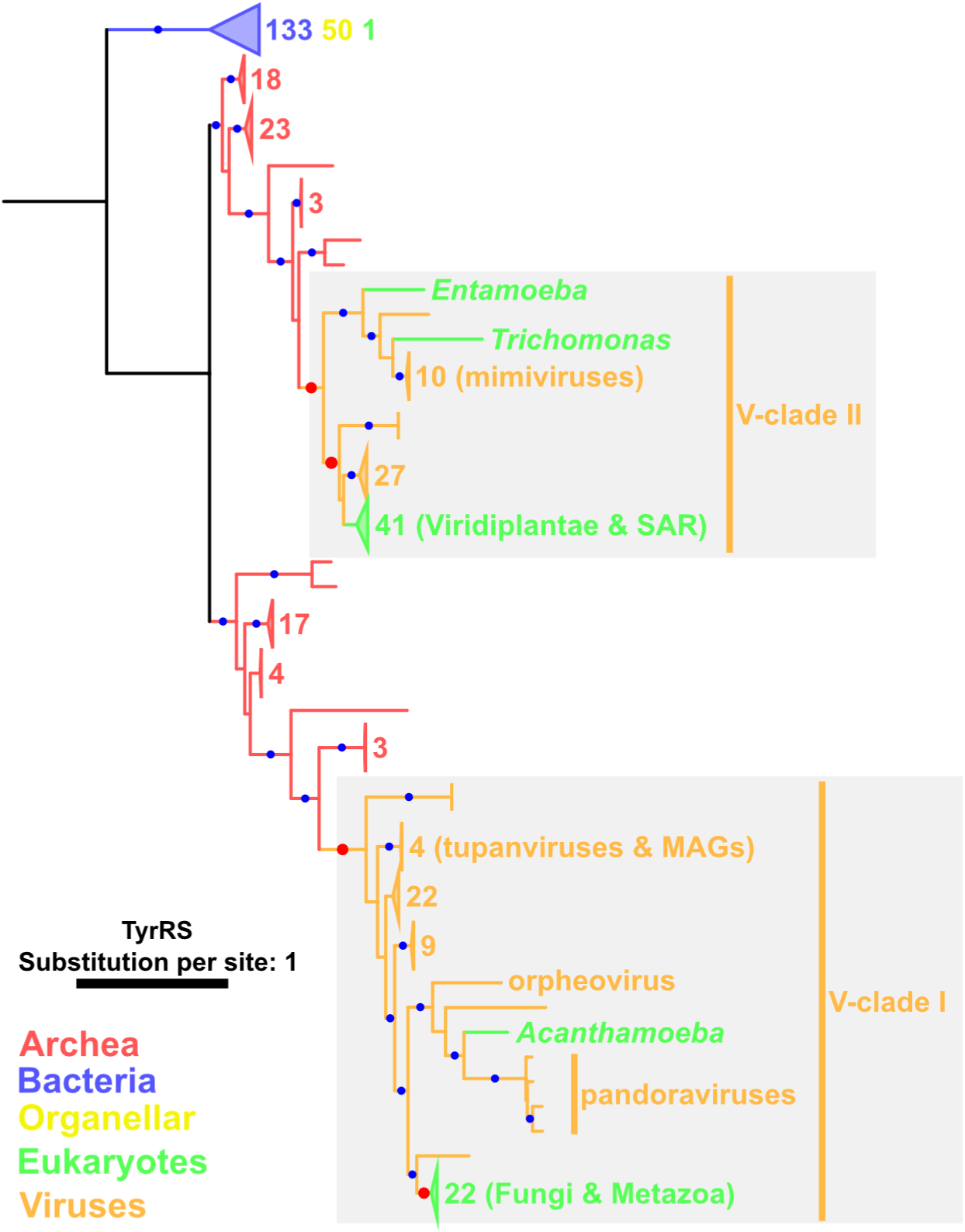
A phylogenetic tree for TyrRS. Red and blue dots indicate significant statistical support (≥80% SH-aLRT & ≥95% UFB). The numbers next to the collapsed clades and their colors indicate the numbers of sequences and their sources, respectively. Red dots and gray area indicate the nodes and the phylogenetic relationship described in the main text, respectively. For eukaryotic sequences, only names of genera or higher taxonomic levels are shown. The substitution model was LG+R9. The root was decided following the work of Furukawa et al. [33].

In the TyrRS tree, most eukaryotic sequences were distributed within two statistically supported clades, both of which were included in viral clades (V-clades I and II; statistically supported) (Fig. 5). From the tree topology, V-clades I and II appear to have originated from Archaea. The eukaryotic clade in V-clade I includes sequences from Fungi and Metazoa and that in V-clade II includes sequences from SAR and Viridiplantae. The Fungi and Metazoa clade in V-clade I was grouped with a clade of orpheovirus and pandoraviruses. Sequences from nucleocytoviruses were also present outside of this group. A parsimonious scenario that explains the nested phylogenetic pattern (i.e., a clade of eukaryotes and viruses surrounded further by other viral sequences) involves an ancient HGT of TyrRS from viruses to eukaryotes. The possibility of a scenario involving multiple acquisitions of TyrRS by viruses from ancient eukaryotes cannot be definitively ruled out.

A similar phylogenetic pattern was observed for V-clade II of the TyrRS tree (Fig. 5). Sequences from SAR and Viridiplantae form a clade (statistically unsupported) within V-clade II. The monophyly of the eukaryotic clade and 29 nucleocytovirus sequences was statistically supported, and this group was further surrounded by other viral sequences (including mimiviruses) to form a larger statistically supported clade. This nested phylogenetic pattern suggests that these eukaryotes acquired TyrRS from nucleocytoviruses. Again, we cannot definitively rule out the possibility of multiple acquisitions by nucleocytoviruses from eukaryotes as two eukaryotic sequences (i.e., *Entamoeba* and *Trichomonas*) were placed in the mimivirus clade.

In terms of the direction of HGT occurring from viruses to eukaryotes, there were several more convincing cases. In V-clade I of the TyrRS tree, *A. castellanii* TyrRS was grouped together with pandoraviruses, indicating an HGT from viruses to *A. castellanii* (Fig. 5). Similar virus-to-eukaryote HGT cases were observed in the trees of SerRS (a *Hydra vulgaris* branch in a viral tree; Fig. S5B) and AsnRS (a *Thalassiosira pseudonana* branch embedded in a large viral tree; Fig. S6A). A *Reticulomyxa filosa* branch in a HisRS tree also suggests a similar HGT event, although this scenario was not statistically supported (Fig. S6B).

### HGT between nucleocytoviruses and prokaryotes (Prok to V)

Nucleocytoviruses exclusively infect eukaryotes, and a majority of detected HGTs were between eukaryotes and nucleocytoviruses. However, our analysis also detected cases of HGTs between viruses and prokaryotes.

A clear case was observed in the tree of ValRS, in which two ValRSs from tupanviruses were grouped together with sequences from *Rickettsia prowazekii* and an organelle of *Ixodes scapularis* (Fig. 6A). This group was further surrounded by bacterial sequences. *R. prowazekii* is an obligate intracellular parasitic bacterium, while *I. scapularis* is a tick species that hosts *Rickettsia* species. These results suggest gene flow from bacteria to nucleocytoviruses and another potential flow from bacteria to ticks. Some *Rickettsia* species are symbionts of amoeba, such as *Acanthamoeba* species, which are known as hosts of tupanviruses [26,35]. Therefore, HGT from symbiont bacteria to viruses within amoeba cells is also a possible source of gene flow. The ValRS tree shows another possible transfer between bacteria and viruses, although the branching position of the viral clade (containing 12 sequences) was not statistically supported (Fig. 6A).

**Figure 6.**
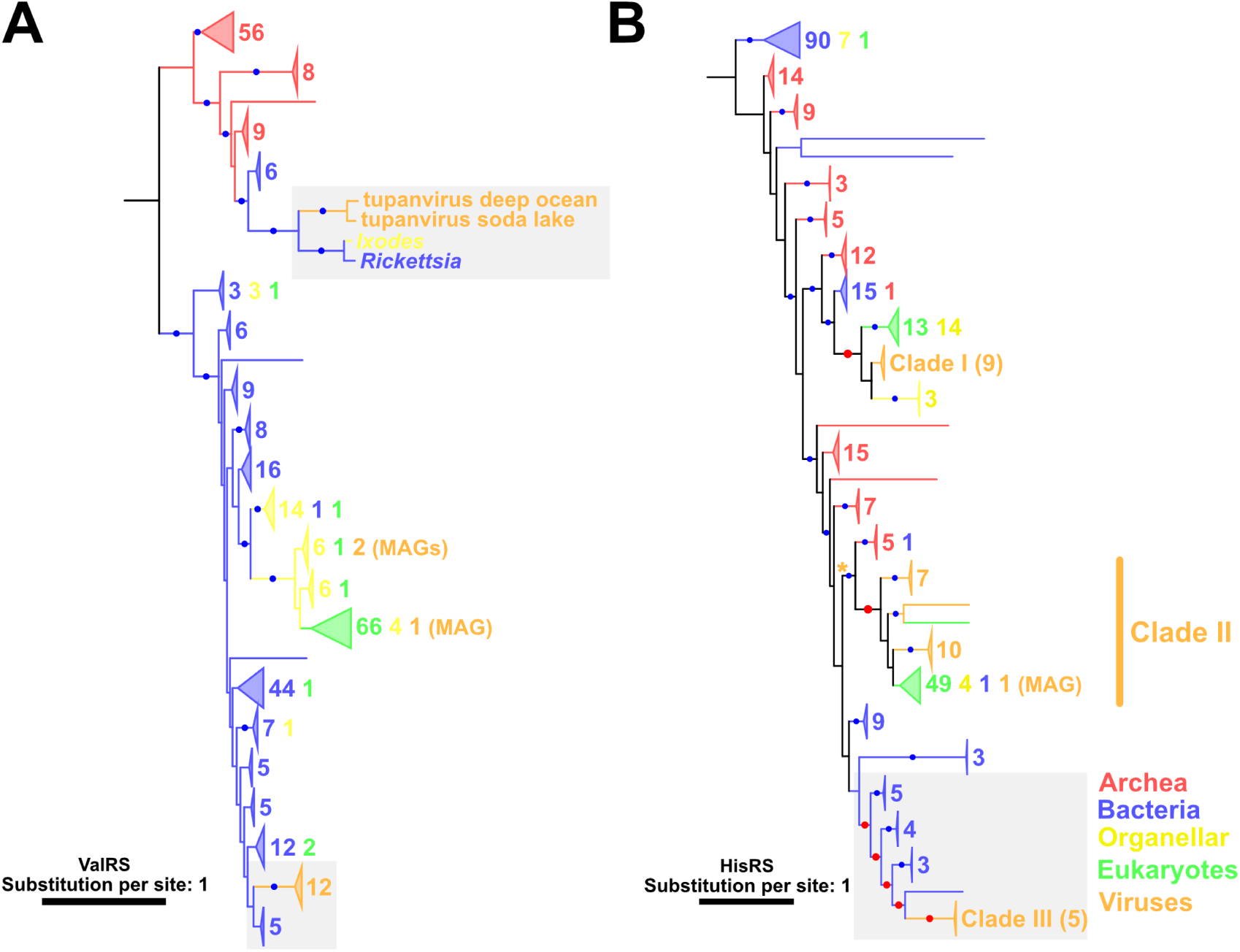
Phylogenetic trees of (A) ValRS and (B) HisRS. Blue dots indicate nodes with significant statistical support (≥80% SH-aLRT & ≥95% UFB). The numbers next to the collapsed clades and their colors indicate the numbers of sequences and their sources, respectively. (B) An asterisk indicates the nodes within which the phylogenetic relationship was further investigated in Figure S6. The substitution models were (A) LG+R10 and (B) LG+R10. The root was decided following the work of Furukawa et al. [33].

Another clear case of HGT from bacteria to nucleocytoviruses was inferred in the HisRS tree. One of the three HisRS clades (i.e., Clade III), which consists of five viral MAGs, was grouped with bacterial sequences with statistical support, indicating the bacterial origin of this viral sequence clade (Fig. S6B). An ancient bacterial origin for viral sequences was also suggested for Clade II of AsnRS, although the true source organisms (bacteria or eukaryotes/organelles) could not be definitively determined (Fig. 3B).

We also noticed that the orpheovirus GlyRS sequence was inside an archaeal clade, suggesting a relatively recent HGT from archaea to nucleocytoviruses (Fig. 2A). Additional possible cases of ancient HGT from archaea to viruses are suggested for TyrRS (Fig. 5; as described above).

### Evolution of aaRSs within nucleocytoviruses

Investigation of aaRSs within viral clades further revealed complex evolutionary pathways of these viral aaRSs. Most of the aaRSs were encoded by *Imitervirales* members, which encode 730 out of 780 aaRSs identified in this study and over half of these aaRSs (n=398) were found in the aaRS-rich clade (Fig. 1). Some aaRSs (10 aaRSs) encoded in this aaRS-rich *Imitervirales* clade appear to have been acquired by the common ancestor of this clade, given the topology of the viral aaRS trees (Fig. S8). In those aaRS trees, clades consisting of sequences from the aaRS-rich *Imitervirales* clade included small numbers of sequences from other *Imitervirales* or viral orders (Fig. S8). These results suggest a scenario in which the common ancestor of the aaRS-rich *Imitervirales* clade acquired these aaRSs from cellular organisms and then spread them to other viruses through virus-to-virus HGTs (hereafter referred to as vHGTs).

In addition to the recent vHGTs, several trees of aaRSs (HisRS, MetRS, ArgRS, and TrpRS) showed the signal of more ancient vHGTs. In these aaRSs, those encoded in the aaRS-rich *Imitervirales* and other *Imitervirales* formed separate clades (Fig. 7). The trees of HisRS and MetRS V-clade I showed statistically supported monophyly of viral aaRSs (Fig. 7A). Within the clades, the separation between the clade for the aaRS-rich *Imitervirales* and that for other *Imitervirales* was statistically supported. In ArgRS and TrpRS, monophyly of the major clade of the aaRS-rich *Imitervirales* was statistically supported and sequences from the other *Imitervirales* were located outside of the clade (Fig. 7B). These results suggest that vHGTs occurred between the common ancestor of the aaRS-rich *Imitevirales* clades and other *Imitervirales*. We acknowledge that an alternative scenario cannot be definitively ruled out, in which the common ancestor of *Imitervirales* acquired the aaRSs and they were subsequently lost in many lineages outside the aaRS-rich clade over the course of evolution.

**Figure 7.**
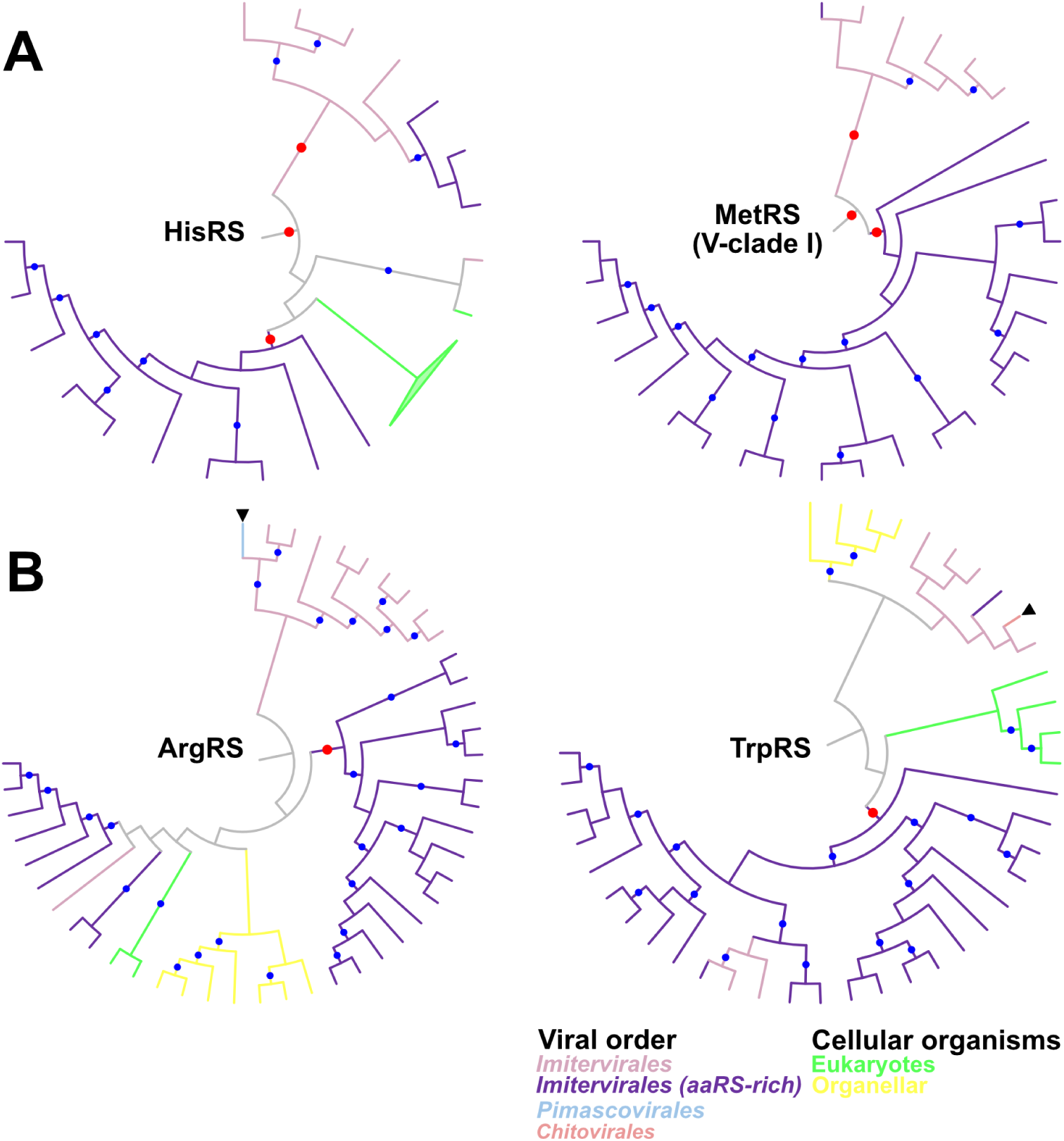
Phylogenetic trees for viral clades found in IleRS, AsnRS, and LysRS. Names of aaRSs are shown in the center of each tree. Monophyletic clades were obtained from the whole phylogenetic trees. Blue and gray dots indicate nodes statistically supported by UFB and SH-aLRT and those supported by TBE, respectively. Node and branch colors indicate the order of nucleocytoviruses. Members of the aaRS-rich clade within Imitervirales are shown in different colors. Black arrowheads indicate putative HGTs between viral orders. Branch length was ignored.

Phylogenetic trees of viral AsnRS, IleRS, and LysRS suggested their acquisition by the common ancestor of *Imitervirales* (Fig. 8). Their phylogenetic trees also showed several signals of vHGTs both within *Imitervirales* and with other viral orders. However, a majority of the sequences from the aaRS-rich *Imitervirales* clade form a relatively large clade inside the whole *Imitervirales* clades, which suggests that the common ancestor of *Imitervirales* acquired the aaRSs (Fig. 8). AsnRSs and IleRSs were identified in the genomes of isolates that are closely related to mimiviruses (i.e., megaviruses and moumouviruses), but absent from those of mimiviruses. As megaviruses and moumouviruses formed a clade with other closely related *Imitervirales* (tupanviruses), relatively recent gene loss occurred in the common ancestor of mimiviruses.

**Figure 8.**
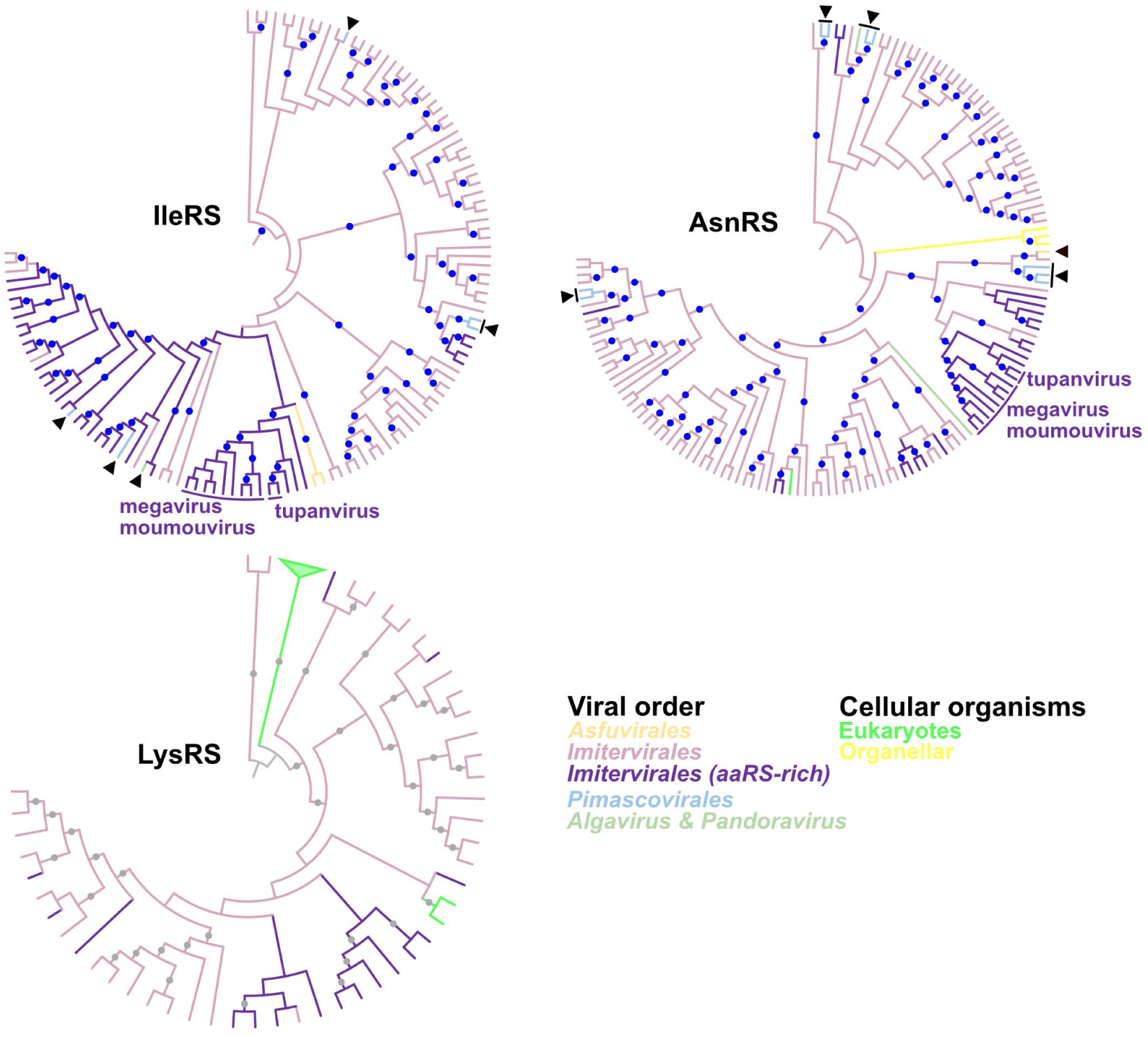
Phylogenetic trees for viral clades found in (A) HisRS and MetRS and (B) ArgRS and TrpRS. Names of aaRSs are shown in the center of each tree. Monophyletic clades were obtained from the whole phylogenetic trees. Blue dots indicate statistically supported nodes. Node and branch colors indicate the order of nucleocytoviruses. Members of the aaRS-rich clade within *Imitervirales* are shown in different colors. Black arrowheads indicate putative HGTs between viral orders. Branch length was ignored.

As reported in cellular organisms [28], some viruses showed displacements of aaRSs by those derived from other species. In the AspRS tree, the topology suggests that AspRS has been vertically inherited at least within the aaRS-rich *Imitervirales* clade (Fig. 9A). However, those encoded in tupanviruses were included in a eukaryotic clade with statistical support.

**Figure 9.**
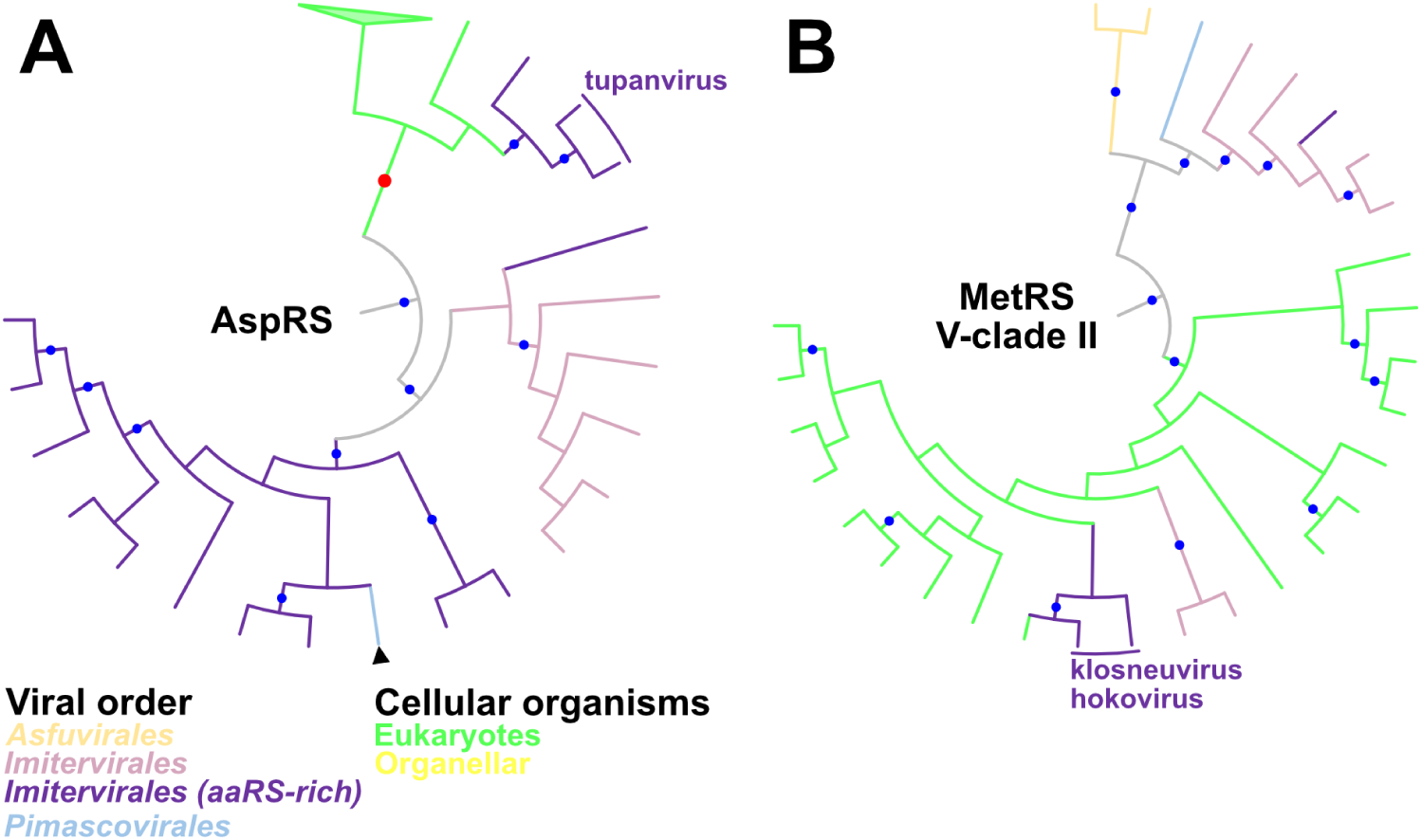
Phylogenetic trees for viral clades found in (A) AspRS and (B) MetRS. Monophyletic clades were obtained from the whole phylogenetic trees. Blue dots indicate statistically supported nodes. Node and branch colors indicate the order of nucleocytoviruses. Members of the aaRS-rich clade within *Imitervirales* are shown in different colors. Black arrowheads indicate putative HGTs between viral orders. Branch length was ignored.

These results suggest that tupanvirus AspRSs were displaced by eukaryotic homologs. Another case was found in MetRS encoded by klosneuvirus and its relative, hokovirus. While MetRSs from the aaRS-rich *Imitervirales* clade formed a major monophyletic clade within V-clade I (Fig. 7), MetRSs from klosneuvirus and hokovirus belonged to a phylogenetically distant eukaryotic clade (Fig. 9B). This suggests that klosneuvirus MetRS was displaced by eukaryotic homologs. Putative gene displacements were also identified in ProRS and TyrRS trees (Fig. S9), which we discuss in Supplementary Information.

## Discussion

The origin and evolution of viral aaRSs have been debated for around two decades since the discovery of APMV [4,13,14,23,25,26]. In the present study, we used an expanded set of nucleocytovirus aaRS sequences, including those from metagenomic data, with a large dataset of cellular aaRSs to construct their phylogenetic trees. The resultant trees recovered viral aaRS clades and their relationship to cellular aaRSs with strong statistical support, most of which was missing in previous studies. On the basis of our updated aaRS phylogeny, recent transfer from eukaryotes to viruses was identified in the trees of AlaRS, MetRS, TrpRS, ArgRS, and SerRS (Figs. 3, 4, and S5). In contrast, the phylogenetic trees of GlyRS, IleRS, PheRS, AspRS, and GluRS provided statistically supported scenarios of ancient transfers between proto-eukaryotes and nucleocytoviruses (Figs. 2 and S2). A similar or more ancient origin was suggested in the TyrRS tree, where eukaryotic clades were nested inside the tree of viral sequences (Fig. 5). Regardless of the direction of HGTs (viruses to proto-eukaryotes or multiple acquisitions by viruses from proto-eukaryotes), the origin of TyrRS can date back to the proto-eukaryotic period. In total, the origins of eight aaRSs date back to the time before LECA and four aaRSs predate the divergence of major eukaryotic lineages (Table 1). These ancient aaRSs outnumber those predicted to have ancient origins in a previous study (only two: GluRS and HisRS) [25]. Although most HGTs appear to occur between viruses and eukaryotes (including proto-eukaryotes) (Table 1), we also identified statistically supported cases for prokaryote-to-virus HGTs for ValRSs, HisRSs, and GlyRS (Figs. 2 and 6). Taken together, our results firmly establish that nucleocytoviruses acquired their aaRSs from diverse sources of cellular organisms across a wide span of time from the proto-eukaryotic period to more recently.

The evolutionary origin of ancestral genomes of nucleocytoviruses is another controversial issue. Some proposed reductive evolution from extinct cellular domains [4,11], while others suggested a model involving accumulation from small ancestral viruses [13,14,16]. The present study showed that some transfers can be traced back to the proto-eukaryotic period, which indicates that the gene pool of ancestral nucleocytoviruses in this period already included an expanded set of aaRSs (Table 1 and Figs. 2 and S2). Notably, however, some aaRSs (e.g., AlaRS, SerRS, and ArgRS) were embedded in eukaryotic clades and likely to have been recently acquired by nucleocytoviruses, suggesting that the set of aaRSs encoded by ancestral nucleocytoviruses was incomplete. Although reductive evolution can partially account for the current distribution of aaRSs in nucleocytoviruses (as we detected gene losses in mimiviruses; Fig. 8), reduction from the complete set of aaRSs is not supported by the current data. A previous study proposed that the ancestral nucleocytoviruses encoded around 40 genes [14]. The expanded set of aaRSs in ancestral nucleocytoviruses as we suggest here does not necessarily contradict this estimate. Instead, it is plausible that this hypothetical small ancestor corresponds to a more ancestral state than the viruses that we consider in this study. We hypothesize that ancestral nucleocytoviruses infecting proto-eukaryotes had already developed complex genomes, as suggested elsewhere [18].

Viruses of the order Imitevirales encode 730 out of 780 nucleocytovirus aaRSs identified in this study, and a clade in this viral order was particularly enriched in aaRSs (Fig. 1). The phylogenetic distribution of viral aaRSs within the viral clades indicates that this aaRS-rich clade provides its viral aaRSs to the other nucleocytoviruses through vHGT (Figs. 7 and S8). We also identified HGTs from members of *Imitevirales* to viruses belonging to other viral orders. These results are consistent with findings in recent studies [21,36,37] and indicate that vHGT is one of the major drivers behind the spread of viral aaRSs among nucleocytoviruses. The phylogenies of some prevalent aaRSs (i.e., IleRS, AsnRS, and LysRS), in contrast, imply vertical evolution of viral aaRSs from the common ancestor of *Imitevirales* (Fig. 8). Moreover, we identified the putative displacement of aaRSs (Figs. 9 and S9). Clear cases were observed in AspRS, MetRS, ProRS, and TyrRS of tupanviruses and klosneuviruses. These viruses have a complete or near-complete set of aaRSs, the origin of which has been debated [25,26]. Our phylogenetic trees indicate that their aaRSs are often displaced by eukaryotic homologs, although their ancestors likely inherited aaRSs from the common ancestor of the aaRS-rich *Imitervirales*. These complex evolutionary trajectories involving HGTs, gene losses, and displacements of viral aaRSs may account, at least in part, for the previous disagreements regarding their origins.

Nucleocytoviruses and eukaryotes evolve through tight interactions via HGTs in both directions: from eukaryotes to viruses and from viruses to eukaryotes [3,20]. Our analyses of aaRSs suggest that most cases are HGTs from eukaryotes to viruses. However, we also identified recent HGTs from viruses to eukaryotes, indicating that the HGTs are not limited to the ancient period but still ongoing evolutionary events. Sequences from *Acanthamoeba castellanii*, *Hydra vulgaris*, *Thalassiosira pseudonana*, and *Reticulomyxa filosa* are embedded in the viral clades in the trees of TyrRS, SerRS, AsnRS, and HisRS, respectively (Figs. 5, S5, and S6). *A. castellanii* is a known host of some nucleocytoviruses, suggesting that viral infection sometimes leads to HGTs from viruses to hosts. *H. vulgaris* harbors an endogenous viral region [38,39]. Recently, giant endogenous viral elements were found in algal species [40] and fungal species [41]. Although the functionality of eukaryotic aaRSs derived from viruses is largely unknown, endogenization of viral genomes may account for some of the HGTs from viruses to eukaryotes.

In cellular organisms, aaRSs exceptionally display a high frequency of HGTs despite their essentiality [28,29]. Nevertheless, their phylogeny is informative to dissect the tree of life [27,28,33]. In the present study, we showed that viral aaRSs have similar complex phylogenetic patterns, namely frequent HGTs, gene losses, and displacements. Meanwhile, by tracing the evolutionary histories of 20 aaRSs, we revealed that the ancestral nucleocytoviruses already encoded expanded sets of aaRSs in the proto-eukaryotic era. These results demonstrate that viral aaRSs are also informative markers to resolve the evolutionary history of nucleocytoviruses and their relationship with cellular organisms. Previous studies aimed at tracing viral evolution mainly focused on functionally conserved genes [1,3,14]. Unlike cellular aaRSs, viral aaRSs do not appear to be essential for viruses. The present study established the presence of auxiliary genes in ancient viruses and their usefulness in resolving the deep evolution of nucleocytoviruses. Analysis of other non-essential viral genes may provide further insights into the properties of ancient viral genomes.

## Materials and Methods

### Sequence dataset

We constructed non-redundant MAGs from 4,168 nucleocytovirus MAGs constructed in previous studies [30–32]. Redundancies among these MAGs were removed as described previously [42]. Briefly, average nucleotide identity (ANI) between MAGs was calculated by dnadiff 1.3 from MUMmer 4.0.0 beta2 [43], and a pair of MAGs that showed ANI greater than 98% with their alignment covering over 25% of the smaller one were clustered together. The largest MAG constructed in *Tara* Oceans Project [32] was selected as the representative of each cluster. If a cluster contained no MAGs constructed in the *Tara* Oceans Project, the largest MAG was selected as representative. In addition, a dataset of 224 reference genomes was obtained from the Global Ocean Eukaryotic Viral database [32]. We used coding regions of sequences predicted in a previous study [32]. Gene annotation was further performed using BLASTP from DIAMOND BLAST v2.0.12.150 [44] against the UniProtKB database (UniProt Consortium 2021) at an E-value threshold of 1e−5.

### Identification of aaRSs

Genes whose best hit was annotated as an aaRS were considered as putative nucleocytovirus aaRSs. Non-functional genes and possible contamination of aaRSs from cellular organisms were excluded from the dataset by using phylogenetic trees constructed for each aaRS of nucleocytoviruses and cellular organisms collected from a previous study [33]. Non-functional genes were manually identified as short sequences and long branch sequences of nucleocytovirus. Decontamination of sequences derived from cellular organisms was performed as follows. We first detected nucleocytovirus aaRSs that were not included in clades that comprise multiple nucleocytovirus aaRSs in each tree. If contigs that encode the detected aaRS sequence were from a MAG and contained no nucleocytovirus core gene or gene whose best hit taxonomy was nucleocytovirus, we removed the aaRSs as contamination from cellular organisms.

### Phylogenetic analysis

Nucleocytovirus eight core genes were identified in the dataset by considering the DIAMOND BLASTP hits that were registered in NCVOG [45]. Multiple sequence alignments were performed by MAFFT v7.487 [46] with the E-INS-i algorithm, which is suitable for the alignment of sequences in which the number of domains is unknown. Subsequently, the sites with gaps in more than 75% of aligned sequences were removed by using trimAl v1.4.rev15 [47]. Phylogenetic trees were constructed within the ML framework by IQ-TREE 2.1.3 [48]. The amino acid substitution model was selected by Model Finder [49]. Bootstrap values were computed by the SH-aLRT method [50] and the Ultrafast bootstrap (UFB) method [51]. Clades with >80% SH-aLRT and >95% UFB were considered to be statistically supported. All phylogenetic trees were visualized by iTOL v6 [52].

The origin of nucleocytovirus aaRS was inferred using phylogenetic trees constructed as described above. We ignored the clades comprising fewer than three nucleocytovirus sequences, to reduce the contamination from nucleocytovirus MAGs contaminated by contigs of cellular organisms. We also constructed phylogenetic trees from subsections of data used for tree reconstruction above. These subsections contain a nucleocytovirus clade and its neighboring cellular branches. In addition to SH-aLRT and UFB, transfer bootstrap expectation (TBE) [53] was calculated. While conventional bootstrap approaches count exactly the same branches in bootstrap trees, TBE takes into account similar branches by using transfer distances [53]. The use of TBE support facilitates inferences about the presence of unstable taxa (single taxa that tend to move in and out of clades) and is considered to be particularly useful for deep branches and large datasets. Here, clades supported with >70% TBE were considered to be statistically supported.

## Acknowledgments

We thank the *Tara* Oceans consortium and the people and sponsors who supported *Tara* Oceans. *Tara* Oceans (including both the *Tara* Oceans and *Tara* Oceans Polar Circle expeditions) would not exist without the leadership of the *Tara* Expeditions Foundation and the continuous support of 23 institutes (https://oceans.taraexpeditions.org). This article is contribution number XXX of *Tara* Oceans. We also thank Edanz (https://jp.edanz.com/ac) for editing a draft of this manuscript. Computation time was provided by the SuperComputer System, Institute for Chemical Research, Kyoto University.

