## Supplementary Information for "Complex genomes of early nucleocytoviruses revealed by ancient origins of viral aminoacyl-tRNA synthetases"

Supplementary Text

- Ancient HGT between proto-eukaryotes and nucleocytoviruses (proto-Euk & V)
- Ancient HGT between eukaryotes and nucleocytoviruses (ancient Euk & V)
- Recent HGT from eukaryotes to nucleocytoviruses (recent Euk & V)
- HGT from nucleocytoviruses to eukaryotes (V to Euk)
- HGT between eukaryotes and nucleocytoviruses for which the timing of transfer is less clear (other Euk & V)
- Evolution of aaRSs within nucleocytoviruses

Supplementary Figures 1–9

Supplementary Tables 1 & 2 (Separate files)

Supplementary Data (Separate file)

**Supplementary Text**

**Ancient HGT between proto-eukaryotes and nucleocytoviruses (proto-Euk & V)**

The phylogenetic tree of AspRS indicates an ancient HGT dating back to the proto-eukaryote era (Fig. S2A). Nucleocytovirus AspRSs were distributed in two supported clades (Clades I and II). Clade I consists of 15 viral AspRSs with eukaryotic and organellar sequences and Clade II includes AspRSs from two tupanviruses and a MAG. Clade II was inside a supported clade containing most of the eukaryotic sequences ("eukaryotic clade"). Grouping of Clade I and the eukaryotic clade was statistically supported, indicating an ancient HGT between the viral (Clade I) and eukaryotic clades. It should be noted that the AspRS from *Parastagonospora nodorum* is an outgroup of the eukaryotic and viral sequences. However, another AspRS encoded by this fungal species is located in a supported fungal clade inside the eukaryotic clade. These results suggest that the sequences inside the fungal clade are vertically inherited, and *P. nodorum* AspRS outside of the eukaryote and viral clade was horizontally acquired. Therefore, we conclude that the root of the eukaryotic clade represents LECA and the HGT between nucleocytovirus and eukaryotic clades occurred in the proto-eukaryotic period.

The GluRS tree exhibited a similar phylogenetic pattern (Fig. S2B). Nucleocytovirus- and eukaryote-derived GluRSs were grouped together with statistical support. Although GluRSs from *Reticulomyxa filosa* and *Naegleria gruberi* were found in an outgroup of the clade of viruses and eukaryotes, these two species also appeared in a large eukaryotic clade. This distribution pattern suggests that the outgroup sequences were horizontally acquired ones, suggesting that the root of the major eukaryotic clade represents LECA. Monophyly of the eukaryotic clade was statistically supported, suggesting an ancient HGT between nucleocytovirus and proto-eukaryotes before LECA.

The topologies of the LysRS and LeuRS trees are also suggestive of HGTs between nucleocytoviruses and proto-eukaryotes (Fig. S2C and D). Monophyletic groupings of eukaryotes and nucleocytoviruses were not supported by UFB or SH-aLRT. However, in a subsection of trees, TBE supported the grouping of eukaryotes, the grouping of nucleocytoviruses, and the grouping of both, suggesting ancient HGTs between viruses and proto-eukaryotes (Fig. S3B and C).

**Ancient HGT between eukaryotes and nucleocytoviruses (ancient Euk & V)**

The AsnRS tree suggests an ancient HGT between eukaryotes and nucleocytoviruses (Fig. 3B). Eukaryotic AsnRS sequences were distributed in two clades, one of which was sister to a clade composed of five nucleocytovirus MAGs (Clade I). The eukaryotic clade and viral Clade I, and their grouping, were statistically supported. The eukaryotic clade encompasses a wide range of taxonomy (*Amorphea*, *Rhodophyta*, *Discoba*, and *Metamonada*), thus indicating an ancient HGT between eukaryotes and viruses before their divergence, although it is unclear if this HGT occurred before LECA.

In the ThrRS tree, Clade I nucleocytovirus ThrRSs were grouped together with many eukaryotic sequences, with the exception of sequences from *Metamonada* species (i.e., *Spironucleus*, *Giardia*, *Trichomonas*), which are located as outgroups (Fig. S4). The grouping of eukaryotes and viral sequences was statistically supported. Inside this group, a sequence from *Paramecium tetraurelia* (Ciliophora) was located as an outgroup of a nucleocytovirus clade. However, sequences from *P. tetraurelia* were also included in a large eukaryotic clade with other Ciliophora species (*Stylonychia lemnae*, *Oxytricha trifallax*, and *Tetrahymena thermophila*), indicating that the outgroup sequence may have been horizontally acquired. Therefore, the root of the eukaryotic clade represents a common ancestor of a wide range of eukaryotes, which are sister to the nucleocytovirus clade. Taken together, nucleocytoviruses probably acquired ThrRS before the divergence of most eukaryotes, although the statistical support was insufficient to draw definitive conclusions about this.

**Recent HGT from eukaryotes to nucleocytoviruses (recent Euk & V)**

The orpheovirus ArgRS was grouped with ArgRSs from Amoebozoa and Fungi species (Fig. S5A). This clade was sister to organellar sequences of Metazoa. Although the precise phylogenetic location of orpheovirus ArgRS could not be determined due to low branch support, this result indicates that orpheovirus acquired its ArgRS from eukaryotes after the establishment of Amorphea (a supergroup including Metazoa, Fungi, and Amoebozoa).

SerRSs from tupanviruses and orpheovirus were distantly related to many MAG-derived viral sequences and located inside a large supported eukaryotic clade, suggesting recent acquisition from eukaryotes (Fig. S5B). However, this eukaryotic clade encompasses a wide variety of taxa, and the precise phylogenetic locations of these viral sequences were unclear because of low statistical support, thus obscuring the precise timing of their acquisition.

Although some ThrRSs (tupanviruses and a MAG) and MetRSs (klosneuvirus, hokovirus, and two MAGs) also suggested recent acquisition from eukaryotes by the viruses, timing and sources were not determined unambiguously due to the limited support (Figs. 3 and S4).

**HGT from nucleocytoviruses to eukaryotes (recent Euk &V)**

HisRS Clade II encompassed some of the eukaryotic sequences (e.g., Metazoa, Fungi, Viridiplantae, Amoebozoa, and Alveolata) (Figs. 6 and S6). Monophyly of this clade was statistically supported. Inside Clade II was a supported nucleocytovirus clade (seven MAGs), which was located as an outgroup of a clade encompassing other viral (10 sequences) and eukaryotic sequences, suggesting the possibility of gene transfer from viruses to eukaryotes. However, as the grouping of the eukaryotes and the 10 viral sequences was not supported, different scenarios are also possible.

**HGT between eukaryotes and nucleocytoviruses for which the timing of transfer is less clear (other Euk & V)**

Nucleocytovirus ArgRSs (Clade II), ProRSs, and GlnRSs were located close to eukaryotic sequences, but limited statistical support hampered the delineation of specific scenarios (Figs. S5A and S7). In addition, the trees of TrpRSs, HisRSs (Clade I), and SerRSs (Clade I) showed a signature of HGTs with eukaryotes, but limited statistical support obscured the timing of gene transfers (Figs. 4, 6, and S5B).

A large number of nucleocytovirus SerRS sequences form a statistically supported monophyletic clade (Clade I; with one exceptional sequence from *Hydra vulgaris* as mentioned in the main text) within a statistically supported eukaryotic clade (Fig. S5B). This suggests that nucleocytoviruses acquired the SerRSs from eukaryotes. Similarly, a large TrpRS clade contains most of the eukaryotic sequences, one small nucleocytovirus clade (Clade I), and large clades with viral MAGs (Clade II) (Fig. 4). Clade II included isolated mimiviruses and its monophyly was statistically supported. However, in both SerRS and TrpRS trees, further relationships between specific eukaryote clades and the viral clade were unclear due to the limited statistical support.

One of the three clades of HisRS (Clade I) was located close to clades of eukaryotic sequences and organellar sequences (Fig. 6). The grouping of these viral and eukaryotic sequences was supported, but further relationships were not determined unambiguously.

**Evolution of aaRSs within nucleocytoviruses**

Viral ProRS phylogeny suggests displacement of viral aaRSs by eukaryotic homologs. Nine out of 16 viral ProRSs were encoded in the aaRS-rich Imitevirales and formed a statistically supported clade (Fig. S9A). This clade included sequences from klosneuvirus, hokovirus, and its relatives. Apart from this clade, some viral ProRSs were encoded within the eukaryotic clades, which included sequences from tupanviruses, suggesting displacement of ProRSs in tupanviruses.

Nucleocytoviruses encode 84 TyrRSs, which were mainly found in Imitevirales genomes and formed two phylogenetically distant clades (Fig. S9B). The sequences from the aaRS-rich Imitervirales clade and those from the other Imitervirales apparently form individual monophyletic clades (Clades I and II). These monophyletic clades included minor clades from the other group, which suggests displacement of TyrRSs.

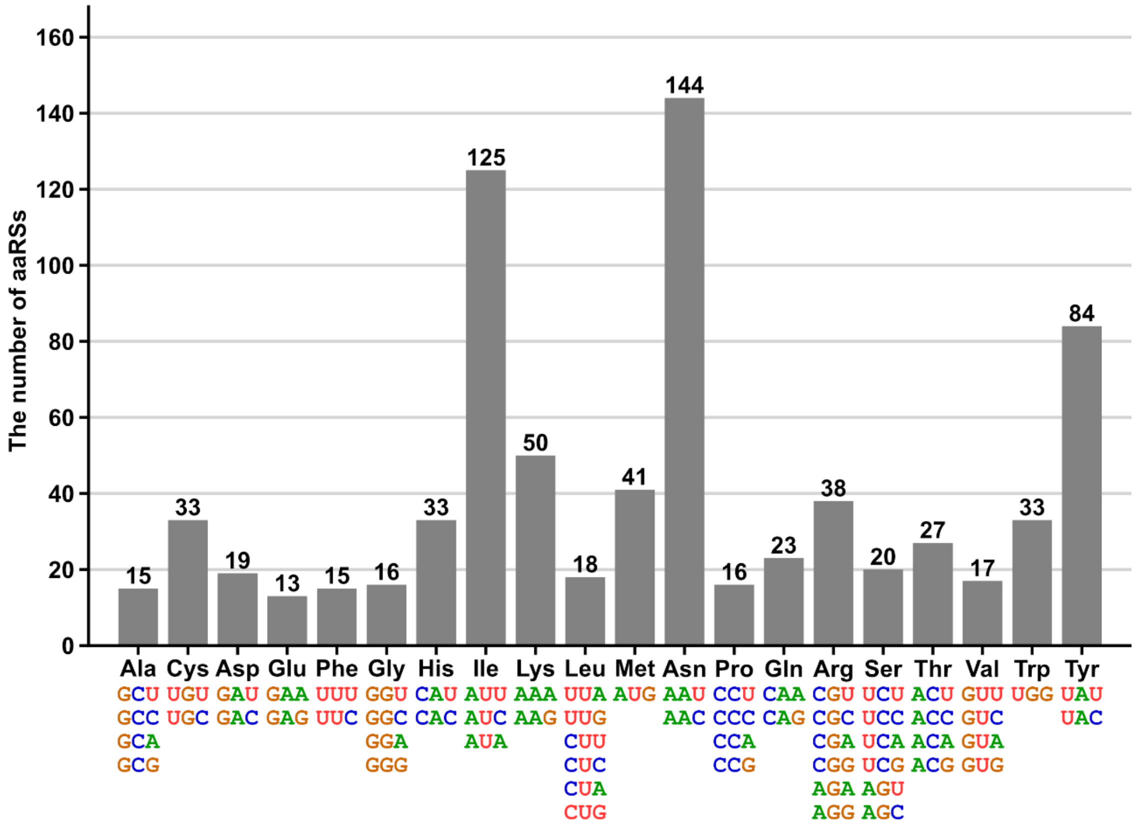


**Figure S1.**

The number of aaRSs detected in nucleocytovirus genomes. The number above each bar indicates the number of each aaRS. Each aaRS is represented as its cognate amino acid. The letters below amino acid names indicate their cognate codons.

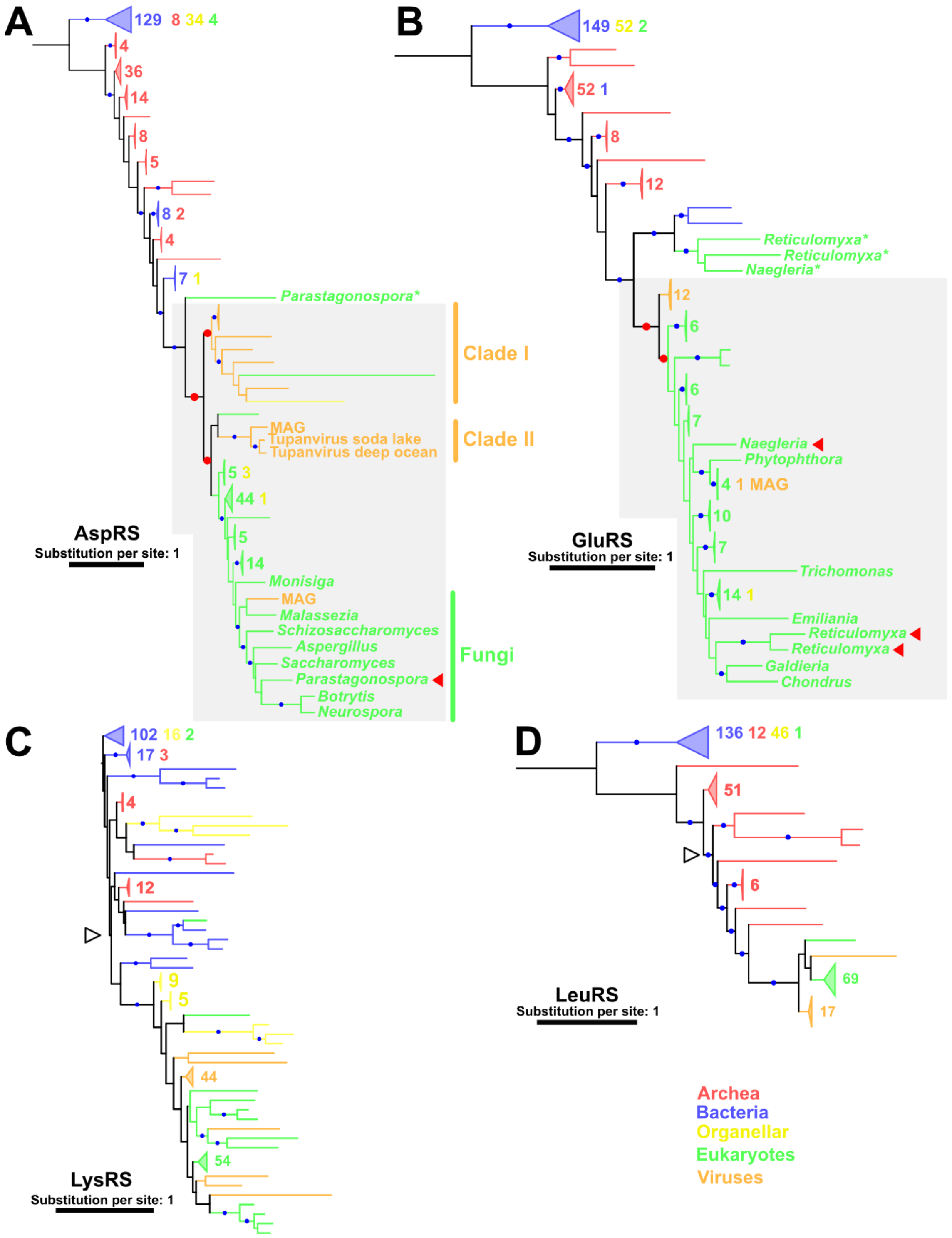


**Figure S2.**

Phylogenetic trees for (A) AspRS, (B) GluRS, (C) LysRS, and (D) LeuRS. Red and blue dots indicate significant statistical support (≥80% SH-aLRT & ≥95% UFB). The numbers next to the collapsed clades indicate the numbers of sequences. Colors of trees and labels represent sources of aaRSs. Red dots and arrowheads represent the supported nodes and the eukaryotic sequences mentioned in the text, respectively. For eukaryotic sequences, only names of genera are shown. Empty arrowheads indicate the node from which subsection trees were built. (A, B) Asterisks indicate the sequences presumably acquired by HGTs and not representing vertical evolution. The substitution models were (A) LG+R10, (B) Q.pfam+F+R10, (C) LG+R10, and (D) LG+R10.

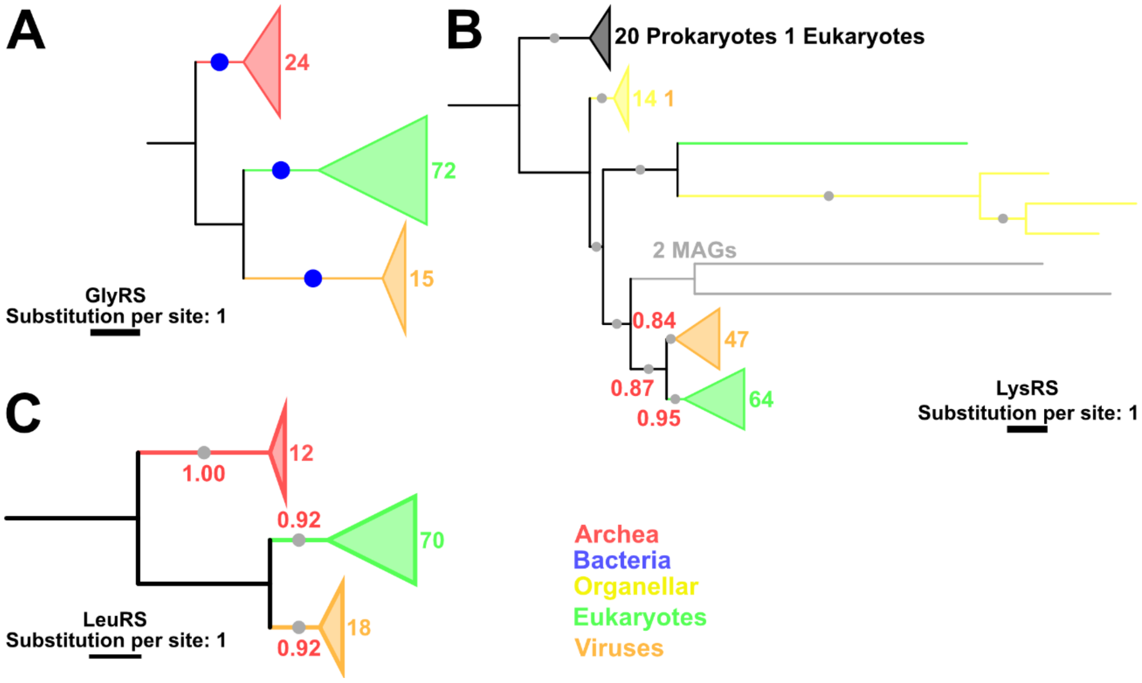


**Figure S3.**

Subsection trees for (A) GlyRS, (B) LysRS, and (C) LeuRS. Blue dots indicate the statistically supported nodes. Gray dots represent nodes supported by TBE. TBE values are shown in red. The numbers next to the collapsed clades and their colors indicate the number of sequences and their sources. The substitution models were (A) LG+I+I+R7, (B) LG+R7, and (C) Q.pfam+F+R7.

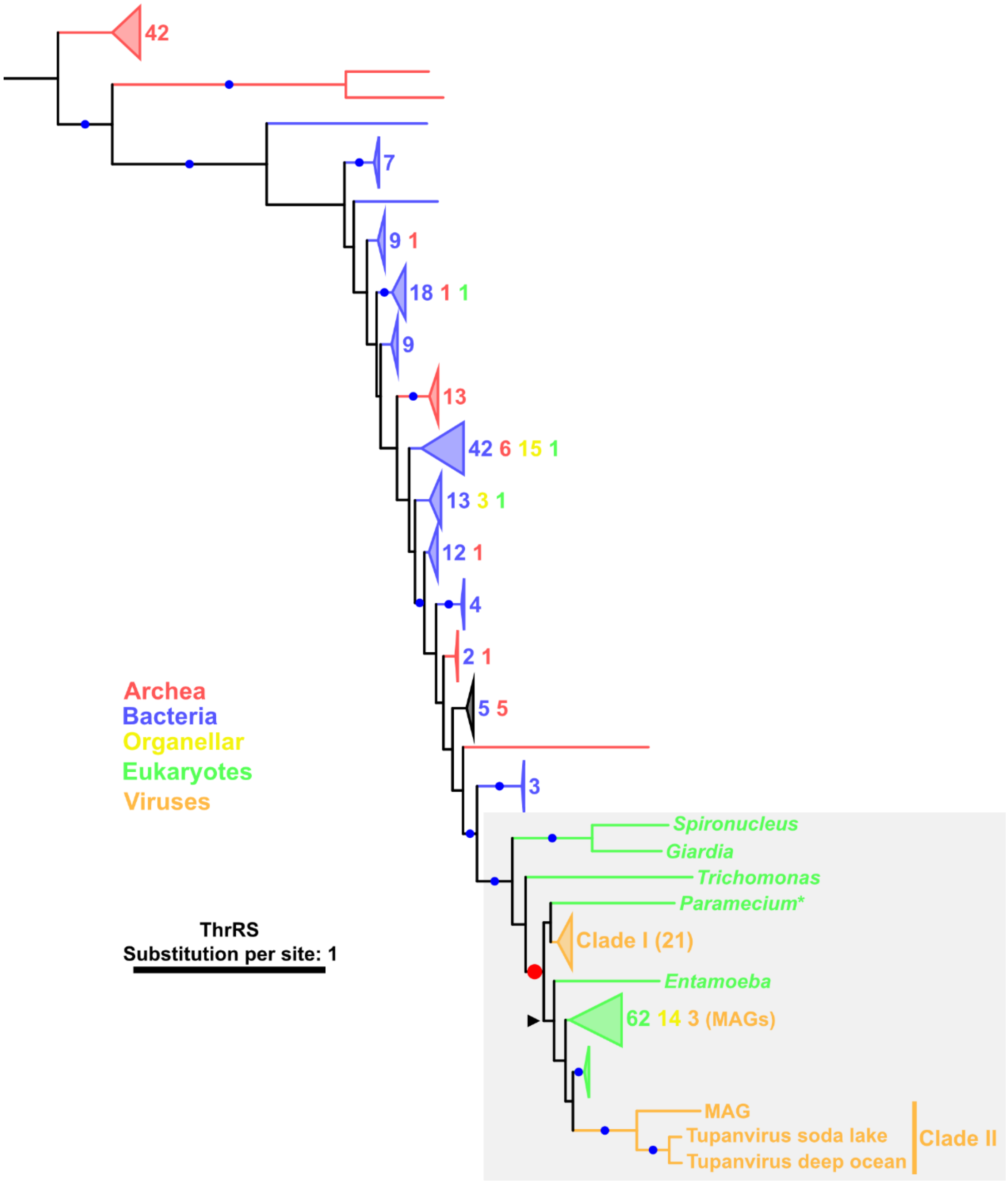


**Figure S4.**

A phylogenetic tree for ThrRS. Red and blue dots indicate significant statistical support. A red dot indicates the node mentioned in the text. Gray area indicates phylogenetic relationships mentioned in the text. The numbers next to the collapsed clades and their colors indicate the numbers of sequences and their sources, respectively. Viral clades are also indicated with the number of sequences in brackets. For eukaryotic sequences, only names of genera are shown. An asterisk indicates the sequence presumably acquired by HGTs and not representing vertical evolution. The substitution model was LG+R9. A black arrowhead indicates the root of the eukaryotic clade mentioned in the text.

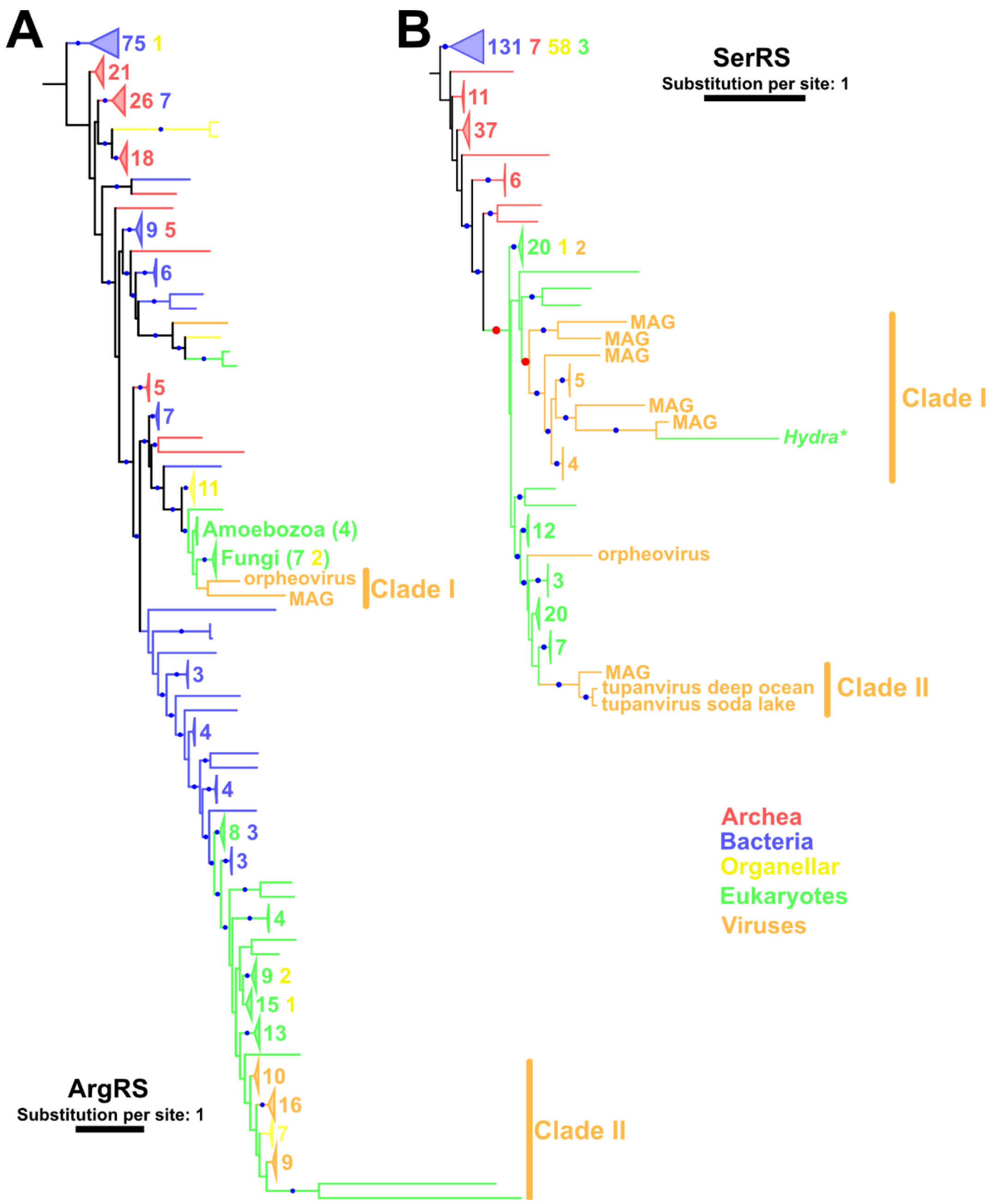


**Figure S5.**

Phylogenetic trees for (A) ArgRS and (B) SerRS. Blue dots indicate the statistically supported nodes. The numbers next to the collapsed clades and their colors indicate the numbers of sequences and their sources. The substitution models were (A) Q.pfam+F+R10 and (B) LG+R9.

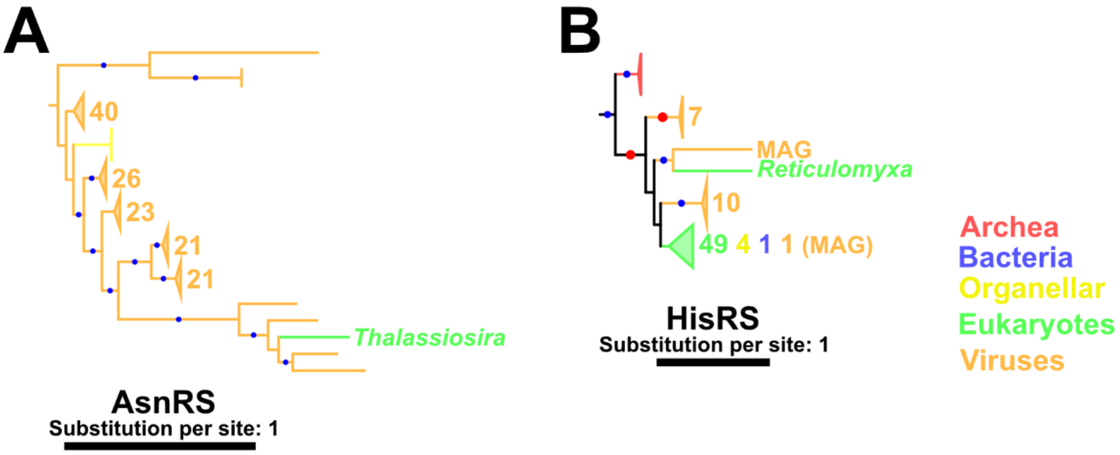


**Figure S6.**

Phylogenetic trees for (A) AsnRS Clade I and (B) subsection of HisRS to highlight eukaryotic sequences within viral clades. Only names of genera are shown for eukaryotic sequences. Blue and red dots indicate the statistically supported nodes and those mentioned in the text. The numbers next to the collapsed clades and their colors indicate the numbers of sequences and their sources. Substitution models were (A) LG+F+R10 and (B) LG+R10 as shown in Figures 3B and 6B, respectively.

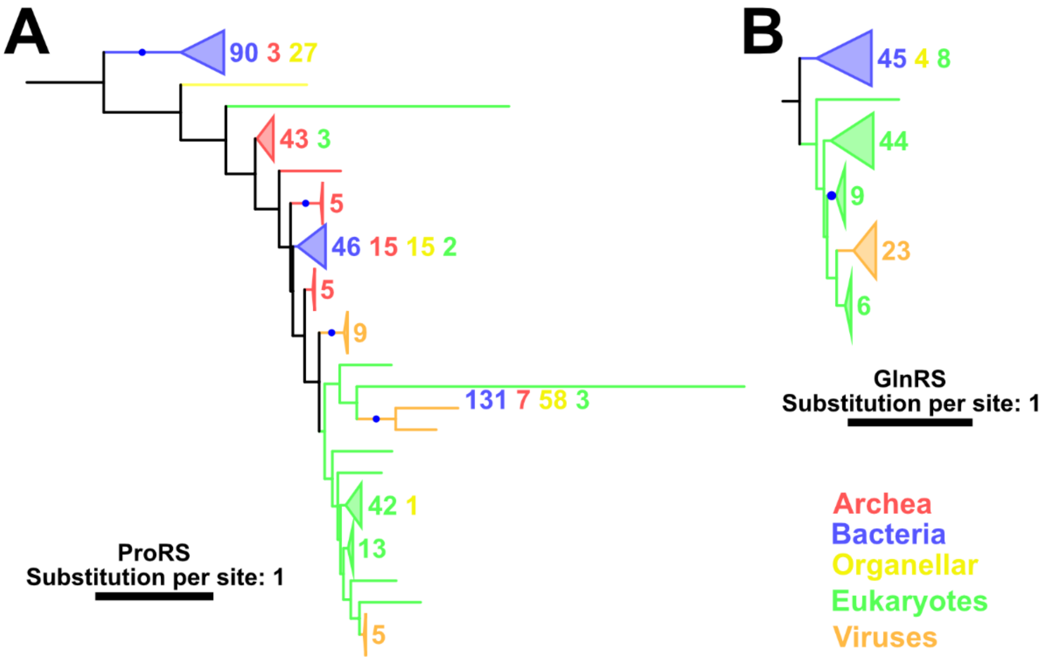


**Figure S7.**

Phylogenetic trees of (A) ProRS and (B) GlnRS. Blue dots indicate the statistically supported nodes. The numbers next to the collapsed clades and their colors indicate the numbers of sequences and their sources. The substitution models were (A) LG+R10 and (B) LG+R8.

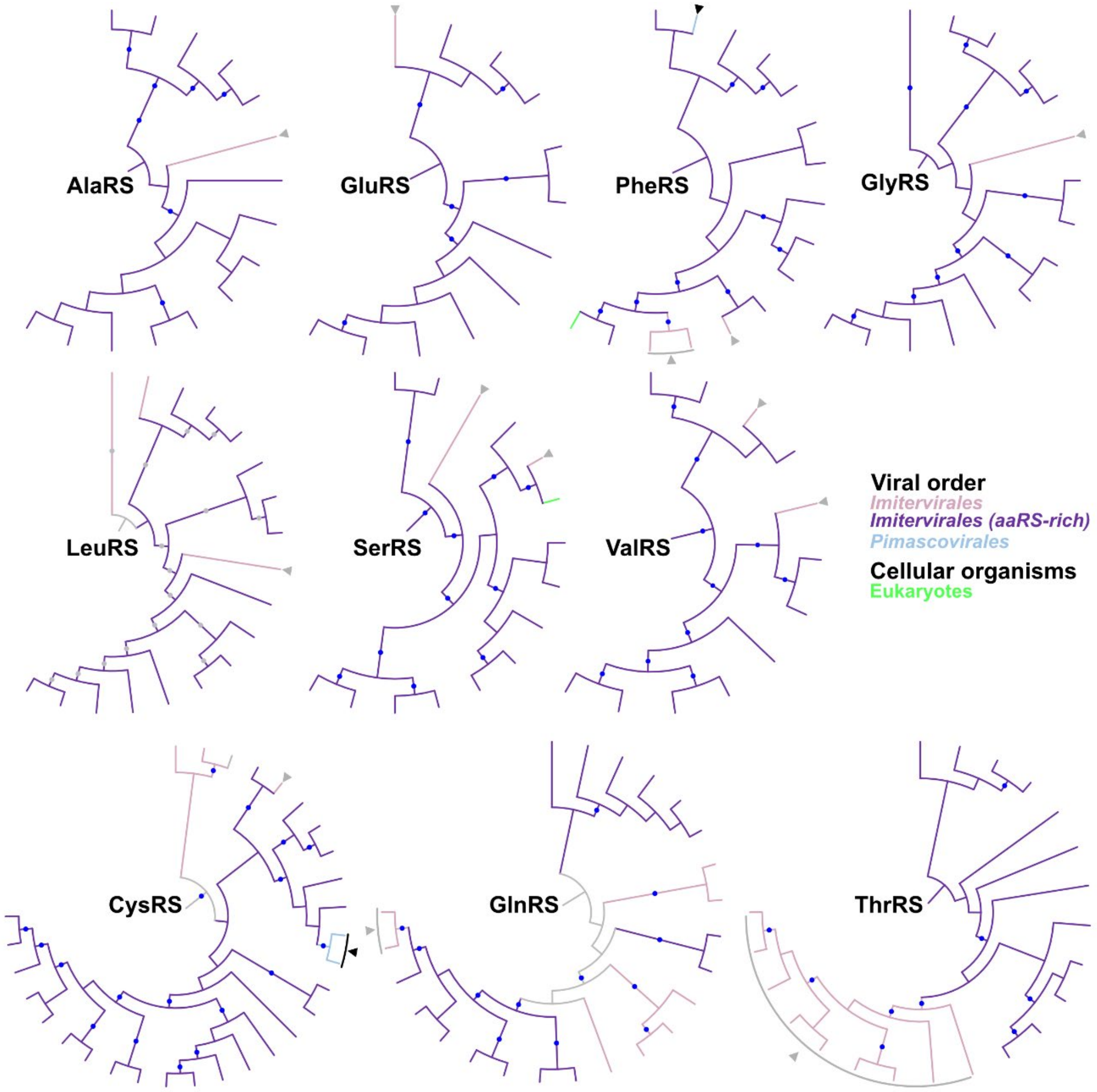


**Figure S8.**

Phylogenetic trees for viral clades. Names of aaRSs are shown in the center of each tree. Monophyletic clades were obtained from the whole phylogenetic trees. Blue and gray dots indicate nodes statistically supported by UFB and SH-aLRT and those supported by TBE, respectively. Node and branch colors indicate the order of nucleocytoviruses or domains of cellular organisms. Members of the aaRS-rich clade within Imitervirales are shown in different colors. Black and gray arrowheads indicate putative HGTs between viral orders and within Imitervirales, respectively. Branch length was ignored.

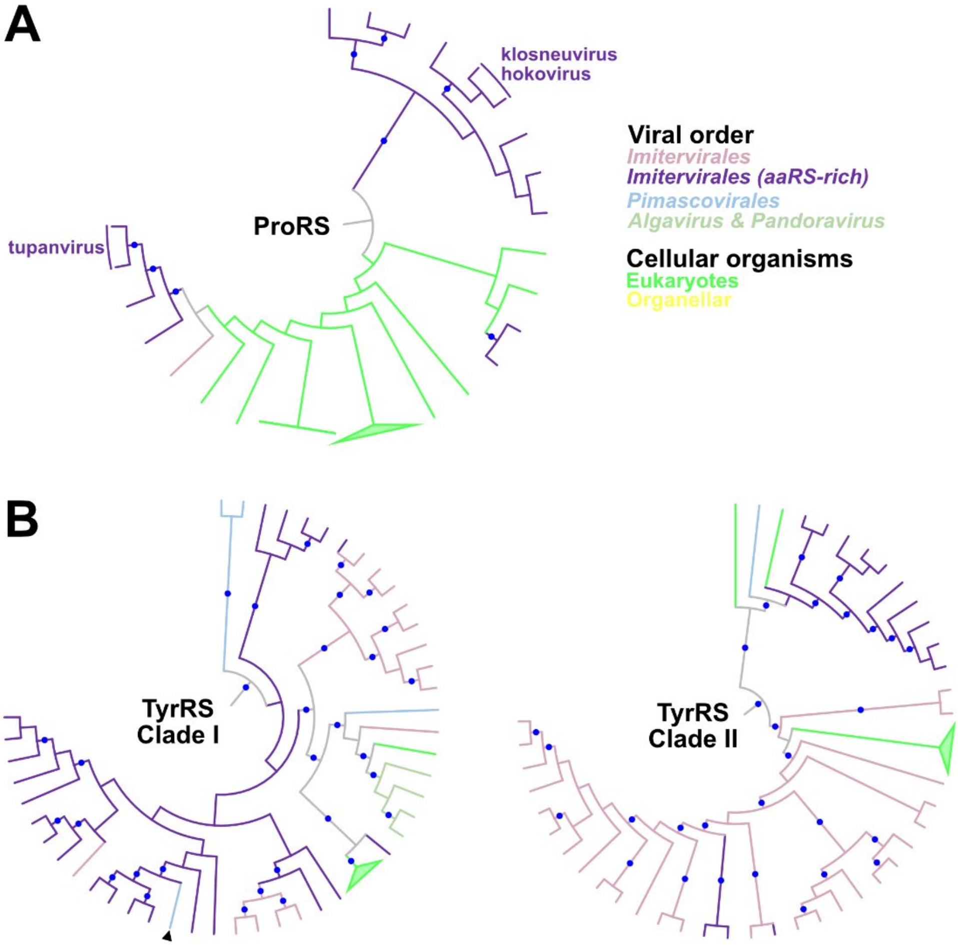


**Figure S9.**

Phylogenetic trees for viral clades of (A) ProRS and (B) TyrRS. The monophyletic clades were obtained from the whole phylogenetic trees. Blue dots indicate statistically supported nodes. Node and branch colors indicate the order of nucleocytoviruses or domains of cellular organisms. Members of the aaRS-rich clade within Imitervirales are shown in different colors. Black arrowhead indicates putative HGT between viral orders. Branch length was ignored.
